# Locomotor deficits in ALS mice are paralleled by loss of V1-interneuron-connections onto fast motor neurons

**DOI:** 10.1101/2020.06.23.166389

**Authors:** Ilary Allodi, Roser Montañana-Rosell, Raghavendra Selvan, Peter Löw, Ole Kiehn

**Author notes:** Corresponding authors: Dr. Ilary Allodi, Prof. Ole Kiehn. These authors contributed equally to the work (Ilary Allodi, Roser Montañana-Rosell).

## Abstract

ALS is characterized by progressive inability to execute movements. Motor neurons innervating fast-twitch muscle fibers exhibit preferential degeneration. The reason for differential vulnerability of fast motor neurons, and its consequence on motor output is not known. Here, we show that fast motor neurons receive more inhibitory synaptic inputs than slow motor neurons, and loss of inhibitory synapses onto fast motor neurons precedes disease progression in the *SOD1*^*G93A*^ mouse model of ALS. Loss of inhibitory synapses on fast motor neurons is accounted for by a loss of synapses from inhibitory V1 spinal interneurons. Deficits in V1-motor neuron connectivity appear prior to motor neuron death and are paralleled by development of specific *SOD1*^*G93A*^ locomotor deficits. These distinct *SOD1*^*G93A*^ locomotor deficits are phenocopied by silencing of inhibitory V1 spinal interneurons in wild-type mice. Silencing inhibitory V1 spinal interneurons does not exacerbate *SOD1*^*G93A*^ locomotor deficits, suggesting phenotypic pathway interaction. Our study identifies a potential cell non-autonomous source of motor neuronal vulnerability in ALS, and links ALS-induced changes in locomotor phenotypes to inhibitory V1 interneurons.

## Introduction

Amyotrophic lateral sclerosis (ALS) is a neurodegenerative disorder characterized by progressive inability to execute movements. During the progression of the disease, somatic motor neurons in the spinal cord and brainstem, as well as corticospinal neurons in the motor cortex, degenerate ^1, 2^. The degeneration of motor neurons is the direct cause of paralysis. Motor neurons that innervate different muscle fibers are not equally affected in ALS. Specifically, motor neurons innervating fast-twitch fatigable fibers – those that produce strong force during movement – are more vulnerable to ALS degeneration than motor neurons that innervate slow-twitch or fast-twitch fatigue-resistant fibers controlling sustained muscle contraction ^2, 3, 4, 5^. Consequently, motor neurons innervating fast-twitch fatigable fibers are first affected in ALS. The reason for this differential vulnerability and its consequence on motor output is unclear ^6, 7, 8^.

While the etiology of most cases of ALS is unknown, mutations in the *SOD1* gene are linked to some familial forms ^9^. In *SOD1* mice ^10^ the SOD1-induced motor neuron degeneration has been linked to motor neuron specific toxicity as well as non-motor neuron-autonomous SOD1-mediated damage, including changes in nearby astrocytes, microglia and oligodendrocytes as well as nerve ensheathing Schwann cells ^11, 12, 13^. In the present study we investigate if the differential vulnerability may be linked to non-motor neuron autonomous circuits in the spinal cord. One such source may be a change in the excitatory or inhibitory synaptic interneuron inputs to motor neurons ^14^. Previous studies have reported inhibitory dysfunction in an ALS *SOD1*^*G93A*^ mouse model, with loss of glycinergic, but not GABAergic, innervation of limb motor neurons ^15, 16, 17^ and reduced glycine receptor in the spinal cord in ALS ^18^. Changes in Renshaw cell-motoneuron connectivity were found in the *SOD1*^*G93A*^ mouse model during disease progression ^19^ and reduction of inhibitory inputs were observed in a *SOD1* zebrafish model ^20^.

Glycinergic synaptic inputs to motor neurons in the spinal cord originate mostly from inhibitory interneurons in the ventral spinal cord, including Ia inhibitory neurons (reciprocal flexor/extensor inhibition), Renshaw cells (recurrent inhibition), and commissural interneurons (crossed inhibition, among other functions) ^21, 22^. The inhibitory interneurons of the ventral spinal cord can be classified into three major classes of neurons characterized by their dorsoventral positioning and expression of developmental transcription factors, with V0 neurons expressing developing brain homeobox protein 1 (Dbx1, commissural), V1 neurons expressing Engrailed 1 (En1), and Renshaw cells and V2b neurons expressing Gata2/3. Selective ablation of these different interneuronal classes leads to specific motor defects: ablation of V0 inhibitory interneurons causes the loss of left-right coordination at slow locomotor speeds ^23, 24, 25^, ablation of V1 interneurons causes a reduction of locomotor speed ^26^, and simultaneous ablation of V1 and V2b neurons eliminates flexor/extensor alternation ^27, 28^.

In the present study, we find 1) motor neurons innervating fast-twitch fatigable fibers receive more glycinergic synapses than motor neurons innervating slow-twitch resistant fibers, 2) a selective loss of glycinergic synapses on fast motor neurons during disease progression in a *SOD1*^*G93A*^ mouse model of ALS, 3) V1, En1-positive, inhibitory interneurons act as the glycinergic population of spinal interneurons accounting for inhibitory synaptic loss during ALS progression, 4) loss of glycinergic and V1 synapses on motor neurons is paralleled by a progressive decrease in locomotor speed which precedes motor neuron degeneration, 5) reversible silencing of spinal V1 neurons phenocopies the *SOD1*^*G93A*^-induced slowing of locomotor speed in wild-type mice, and 6) silencing of V1 neurons has no effect in *SOD1*^*G93A*^ mice, suggesting a phenotypic pathway interaction. These findings implicate a V1 inhibitory interneuron contribution to fast motor neuron vulnerability in ALS and demonstrate that fast motor neuron-specific deficits lead to locomotor slowing prior to motor neuron degeneration in ALS.

## Results

### Fast motor neurons show higher density of inhibitory glycinergic synapses than slow motor neurons

To evaluate the inhibitory connections to fast and slow motor neurons we used a *GlyT2*^*GFP*^ (glycinergic transporter 2) mouse line^29^ where eGFP reliably labels glycinergic neurons and their terminals in the spinal cord^30^. Fast motor neuron somata in the ventral horn were labeled with antibody against Matrix metalloproteinase 9 (MMP9)^8^ (Fig. 1A) while slow motor neurons where labeled with antibody against Estrogen-related receptor beta (ErrBeta)^31^ (Fig. 1C). Only large cells (over 200 mm^2^ in soma area) present in the ventral horn of the spinal cord were quantified, and the synaptic vesicle marker synaptophysin was used to mark putative synaptic terminals^32^. The GFP (glycinergic) positive terminals around somata were specifically selected, reconstructed and density measurements were normalized for the soma areas (Fig. 1B; D). A minimum of 200 motor neurons from the lumbar segment of the spinal cord were quantified per condition. The overall synaptic density was larger on fast motor neuronal somata than on slow motor neurons (MMP9 = 566±60.75 number of synapses/motor neuron per mouse and ErrBeta = 417.19±17.5 synapses/motor neuron per mouse; n = 9; *t test* P= 0.0317) (Fig. 1E), and glycinergic terminals were also found at higher density around fast motor neuron somata (MMP9 = 274.4±29.25 synapses/motor neuron per mouse and ErrBeta = 142.75±39.77 synapses/motor neuron per mouse; n = 9; *t test* P= 0.0008) (Fig. 1F).

**Figure 1.**
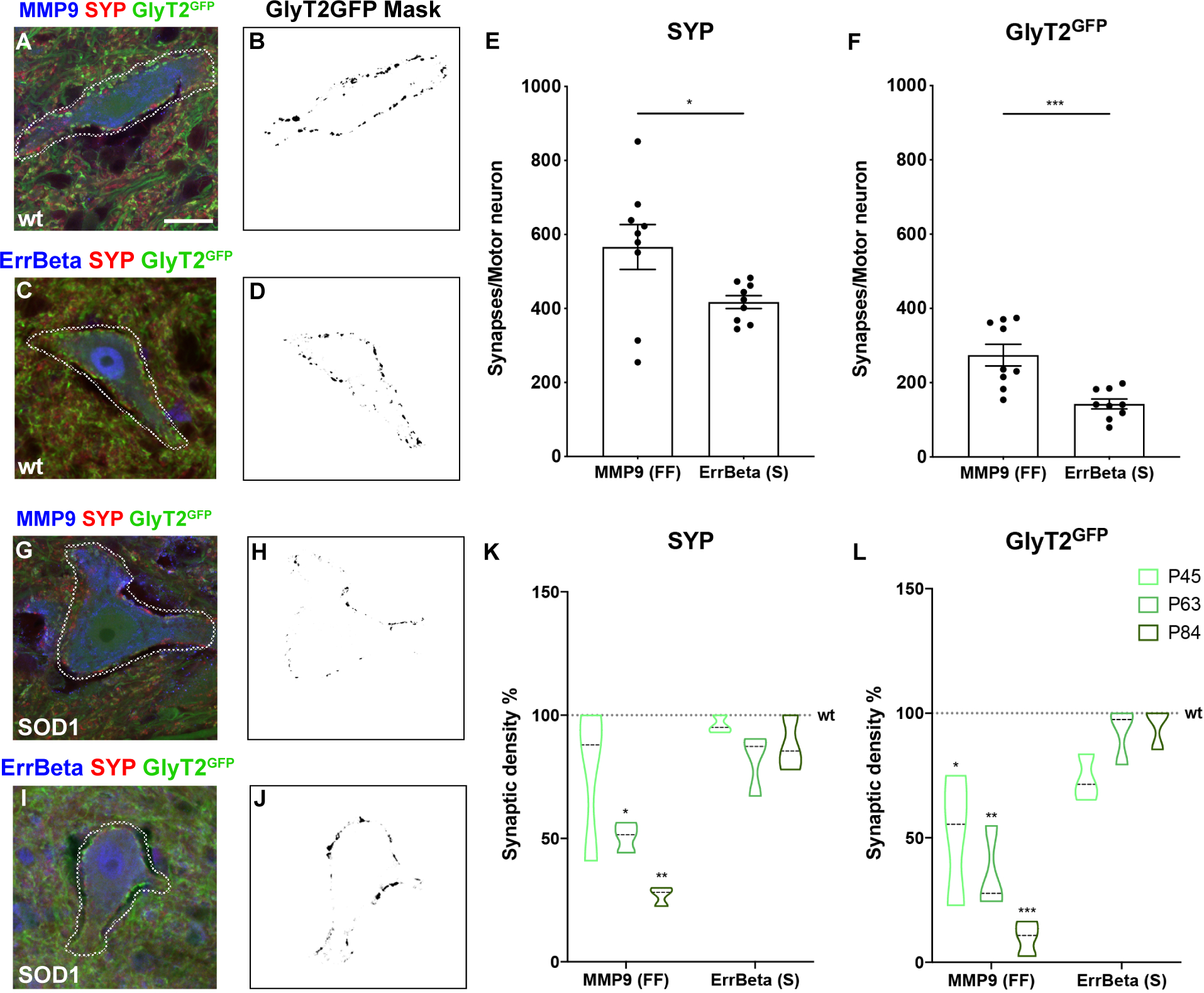
Preferential glycinergic innervation of motor neurons innervating fast-twitch fatigable muscle fibers and loss in a SOD1G93A mouse model. Microphotographs depicting glycinergic inputs onto MMP9^+^ fast motor neurons (A) and ErrBeta^+^ slow motor neurons (C) in a *GlyT2*^*GFP*^ mouse. Synaptic inputs were selected as shown by the dashed line and reconstructed for synaptic density quantifications. Masks in (B) and (D) show processed images. Quantifications expressed as average of synapses per motor neuron – corrected for motor neuron area – of each mouse, show higher synaptic density in MMP9^+^ motor neurons for both synaptophysin (SYP) (MMP9 = 566±60.75 and ErrBeta = 417.19±17.5; *t test* P= 0.0317; n = 9) and GlyT2^GFP^ (MMP9 = 274.4±29.25 and ErrBeta = 142.75±39.77; *t test* P= 0.0008; n = 9) markers. Innervation of fast (G) and slow (I) motor neurons in *GlyT2*^*GFP*^ mice crossed with *SOD1*^*G93A*^ mice at postnatal day (P) 63 and their respective synaptic density reconstruction (H – J). Quantifications of synaptic density (average number of synapses per motor neuron of each mouse) were performed at P45, P63 and P84 for SYP (K) and GlyT2^GFP^ (L) markers in *SOD1*^G93A^ and normalized to age-matching *wt* littermates. MMP9^+^ motor neurons showed progressive reduction of SYP^+^ (one-way ANOVA and Dunnett’s *post hoc*, P45 P= 0.6992; P63 P= 0.0382; P84 P= 0.0027; n = 3 per time point) and GlyT2^GFP+^ inputs (one-way ANOVA and Dunnett’s *post hoc*, P45 P= 0.0121; P63 P= 0.0015; P84 P< 0.0001; n = 3 per time point) when compared with ErrBeta^+^ motor neurons (SYP+ one-way ANOVA and Dunnett’s *post hoc*, P45 P= 0.7919; P63 = 0.1285; P84 = 0.9495; n = 3 per time point) (GlyT2^GFP+^ one-way ANOVA and Dunnett’s *post hoc*, P45 P= 0.057; P63 = 0.9999; P84 = 0.9694; n = 3 per time point). MMP9 (A; G) or ErrBeta (C; I) in blue, SYP in red and GlyT2^GFP^ in green. Scale bar = 10 mm.

### Fast motor neurons lose inhibitory glycinergic connections before slow motor neurons

We next crossed *Glyt2*^*GFP*^ mice with *SOD1*^*G93A*^ mice and performed measurements of synaptophysin (SYP) and eGFP density on MMP9^+^ and ErrBeta^+^ motor neurons (Fig. 1G– I). The phenotype and the genotype of the newly derived *SOD1*^*G93A*^; *Glyt2*^*GFP*^ mice was assessed for copy number of the human *SOD1*^*G93A*^ mutated transgene to verify that copy number was maintained between *SOD1*^*G93A*^; *Glyt2*^*GFP*^ and pure *SOD1*^*G93A*^ mice (S Fig. 1A). The change in weight in the two strains (S Fig. 1B) as well as the survival end point were similar (*SOD1*^*G93A*^; *Glyt2*^*GFP*^ survival = 163.6±14 days n = 4; *SOD1*^*G93A*^ survival = 157.9±10 days n = 7). Moreover, motor neuron number (S Fig. 1C–E) was comparable to the original *SOD1*^*G93A*^ mouse model^33, 34^. Thus, the *SOD1*^*G93A*^; *Glyt2*^*GFP*^ mice exhibited no differences from the original *SOD1*^*G93A*^ strain with respect to ALS phenotype. We analyzed the synaptic density at three time points, postnatal (P) day 45, 63 and 84 and age-matching *wt*; *Glyt2*^*GFP*^ littermates were used as control. A minimum of 200 cells was quantified per motor neuron type (slow vs fast). At postnatal (P) day 63 we found that there was a reduction to around 50 % of synaptophysin inputs (one-way ANOVA P= 0.0382; n = 3) (Fig. 1K) onto MMP9^+^ motor neurons, while eGFP positive inputs were already reduced to 51 % at P45, an earlier timepoint, when compared to wild-type mice (one-way ANOVA P= 0.0121; n = 3) (Fig. 1L). This reduction onto MMP9^+^ motor neurons continued dramatically with ALS progression and was down to 9.94 % at P84 compared to age matched wild-type controls (one-way ANOVA P < 0.0001; n = 3) (Fig. 1L). Moreover, the reduction of eGFP^+^ terminals on fast motor neurons was not due to loss of motor neurons or shrinkage in their size (S Fig. 1C–E). Overall, the general synaptic loss on MMP9^+^ motor neurons – as measured with synaptophysin immunoreactivity – was less than the reduction in glycinergic terminals, suggesting that glycinergic synapses were primarily affected (Fig. 1K). In contrast to these changes, the analysis conducted on ErrBeta^+^ motor neurons showed that glycinergic inputs on slow motor neurons are intact at P84 and comparable to what is seen in age-matched wild-type littermates (Fig. 1K–L).

Together these data demonstrate that MMP9^+^ neurons receive more inhibitory inputs than ErrBeta^+^ motor neurons and that there is a specific soma-near synaptic loss of inhibitory synapses on MMP9^+^ motor neurons that starts before the somatic spinal motor neurons are affected in *SOD1*^*G93A*^ induced ALS.

### Engrailed 1 inhibitory spinal interneurons have more synaptic inputs on fast motor neurons than slow and show selective loss in SOD1^G93A^ induced degeneration

We next tested if the V1 subpopulation of spinal inhibitory neurons was involved in the differential innervation of MMP9^+^ and ErrBeta^+^ motor neurons and in the degeneration-induced synaptic stripping. The Engrailed 1 (En1) transcription factor is found in ipsilaterally projecting neurons in the intermediate and ventral parts of the spinal cord which are known to project to motor neurons innervating the limbs with soma-near connections. These interneurons account for between 39–55 % of inhibitory synapses found on motor neuron somata^27^ and, in the adult mouse, 80 % of these synapses are GlyT2 positive^35^. To visualize the En1^+^ synaptic contacts on motor neurons we first crossed *En1*^*cre*^ ^36, 37^ with the *Rosa26*^*YFP*^ reporter mice^38^. However, the YFP^+^ was not strongly expressed in synaptic terminals (S Fig. 2A– B). We therefore used a viral approach to visualize the terminals of En1^+^-lineage neurons and cell bodies by injecting AAV1-phSyn1(S)-FLEX-tdTomato-T2A-SypEGFP-WPRE^39^ (Fig. 2A) on each side of the lumbar (L) spinal cord at the L2–L3 level (two injections of 80 nl) in P42 *En1*^*cre*^ mice^36, 37^ and P42 *En1*^*cre*^ mice crossed with *SOD1*^*G93A*^ mice. The genotype and phenotype of *SOD1*^*G93A*^;*En1*^*cre*^ did not differ from the non-crossed *SOD1*^*G93A*^ strain, and *SOD1*^*G93A*^;*En1-Cre* mice reached the same end-stage at 158.5±11 (n = 5) as *SOD1*^*G93A*^ alone (S Fig. 2C–D). In *En1*^*cre*^ mice at P63 we found that the En1^+^ soma-near synaptic terminal density (EGFP) was significantly larger on fast motor neurons than on slow motor neurons (MMP9 = 49.2±11.4 and ErrBeta = 33.2±9 synapses/motor neuron per mouse; n = 5; *t test* P= 0.0401 – a minimum of 300 motor neurons were analyzed per condition) (Fig. 2B). In *SOD1*^*G93A*^*-En1*^*cre*^ mice at P63 the overall soma-near synaptic terminal density was visibly reduced as compared to the *wt;En1*^*cre*^ littermates (Fig. 2C–D). Moreover, the number of Tdt positive (En1) neurons was significantly lower in the *SOD1*^*G93A*^;*En1*^*cre*^ mice than in *wt;*En1^cre^ littermates (*wt*;*En-1*^*cre*^ = 99±11.62 %; *SOD1*^*G93A*^;*En1*^*cre*^ = 29.03±13.41 %; n = 5; *t test* P= 0.0040). The synaptic density analysis showed overall loss of soma-near En1^+^ terminals on motor neurons in the *SOD1*^*G93A*^;*En1*^*cre*^ mice at P63 (Fig. 2F–O), with a dramatic reduction to 13 % on fast motor neurons (*t test* P< 0.0001; n = 5) (Fig. 2F–J) and to 53 % on the slow motor neurons compared with *wt*;*En1*^*cre*^ mice (*t test* P= 0.0245; n = 5) (Fig. 2K–O). These data show that inhibitory En1 interneurons dominantly innervate MMP9^+^ fast motor neurons and that these synapses are retracted during the disease progression in *SOD1*^*G93A*^ mice.

**Figure 2.**
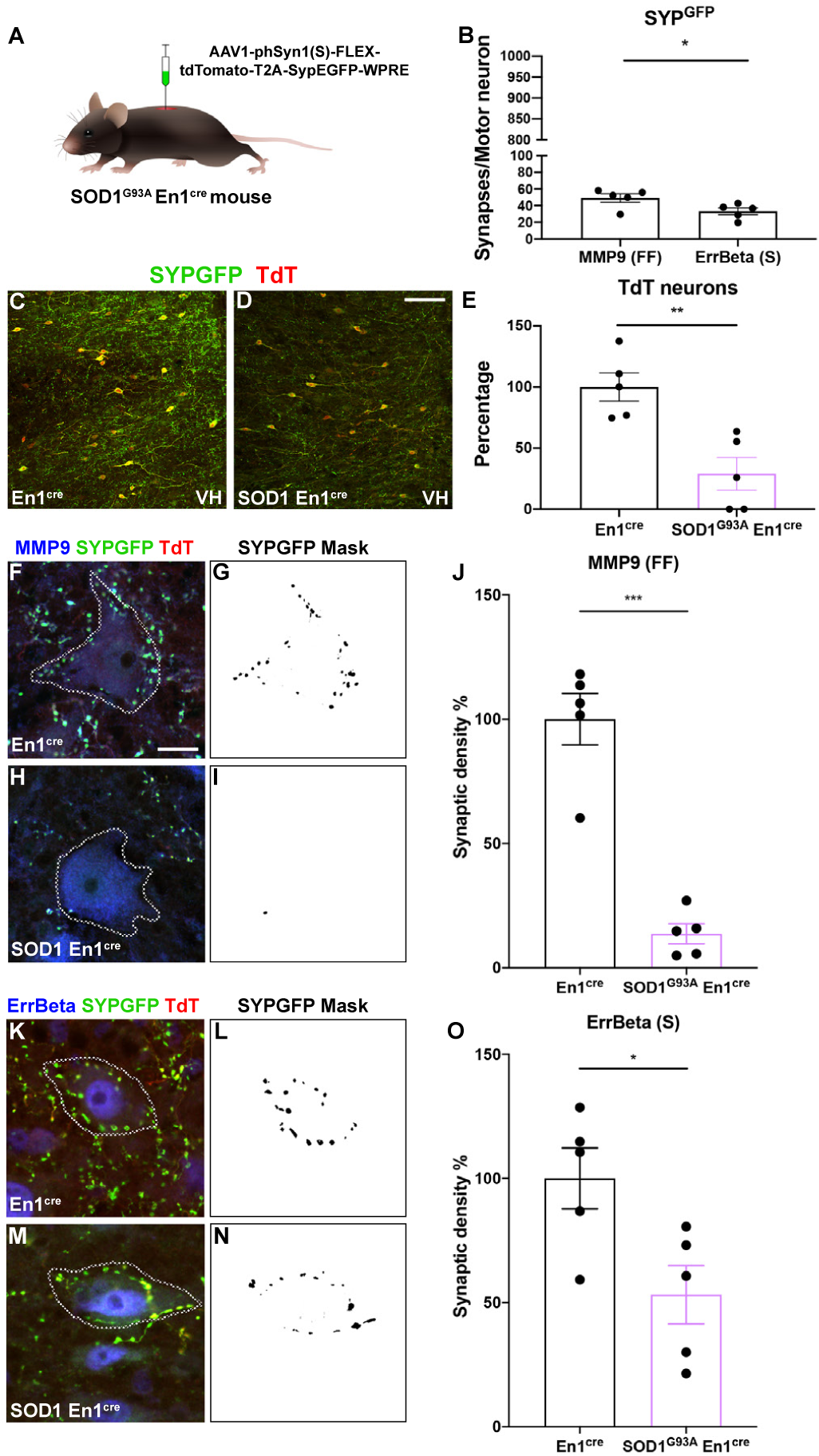
Fast motor neurons receive stronger V1 interneuron innervation than slow motor neurons and V1 synapses are withdrawn in the *SOD1*^*G93A*^ mouse model. (A) Experimental approach: an AAV-phSyn1(S)-FLEX-tdTomato-T2A-SypEGFP-WPRE virus was delivered by intraspinal injections in *En1*^*cre*^ or *SOD1*^*G93A*^;*En1*^*cre*^ mice at P42. (B) In *En1*^*cre*^ mice, MMP9^+^ fast motor neurons were found to receive more inputs than ErrBeta^+^ slow motor neurons, quantifications show average number of synapses per motor neuron in each mouse (MMP9 = 49.2±11.4 and ErrBeta = 33.2±9; *t test* P= 0.0401; n = 5). Microphotographs depicting transduced spinal cords in *En1*^*cre*^ – control condition – (C) and in *SOD1*^*G93A*^;*En1*^*cre*^ animals (D). Synaptic terminals are GFP^*+*^ while *En1*^*cre*^ interneurons are TdTomato+. (E) Percentage of TdT positive interneurons three weeks after viral delivery, number of positive neurons is significantly lower in *SOD1*^*G93A*^;*En1*^*cre*^ than in *En1*^*cre*^ mice (*En1*^*cre*^ = 99±11.62 %; *En1*^*cre*^*-SOD1*^*G93A*^ = 29.03±13.41 %; *t test* P= 0.0040; n = 5 per condition). Synaptic density analysis performed on fast motor neurons in *En1*^*cre*^ (F) and *SOD1*^*G93A*^;*En1*^*cre*^ mice (H), masks are shown respectively in (G) and (I). Percentage of synaptic density (average number of synapses per motor neuron of each mouse) (J) in MMP9^+^ motor neurons revealed a dramatic reduction in GFP^+^ terminals in *SOD1*^*G93A*^ compared to control littermates (*t test* P< 0.0001; n = 5). ErrBeta^+^ motor neurons were analyzed in *En1*^*cre*^ (K) and *SOD1*^*G93A*^;*En1*^*cre*^ *mice* (M), synaptic density was reconstructed (L – N) and quantified (O). Percentage of synaptic density in slow motor neurons showed also a reduction in GFP^+^ terminals in ALS mice (*t test* P= 0.0245; n = 5). TdTomato in red, GFP in green and MMP9 (F; H) or ErrBeta (K; M) in blue. Scale bar in (D) = 100 mm, in (F) = 20 mm.

### *SOD1*^*G93A*^ mice develop progressive slowing of locomotor speed at early time-points while grip strength force is not reduced

Since fast and slow motor neurons are recruited in different motor tasks – with slow motor neurons recruited at low task forces and fast motor neurons at higher task forces – we tested the locomotor performance of *SOD1*^*G93A*^ mice at different postnatal times. Mice were tested weekly between P49 until P112 on a treadmill at a speed of 20 cm/s for 10 s in 3 consecutive sessions (corresponding to fast walk or trot ^24^). Videos were tracked by DeepLabCut analysis ^40^ (Fig. 3A). Over time, *SOD1*^*G93A*^ mice showed progressive inability of coping with the 20 cm/s speed and increased dragging events (Fig. 3B, video 2). To compensate for the reduced locomotor capability developing over time, the speed of the belt was decreased to either 15, 10 or 5 cm/s depending on the severity of the phenotype. The timepoint in which a *SOD1*^*G93A*^ mouse could not sustain a speed of 20 cm/s was called ‘*Onset of locomotor phenotype’*. Notably, between P49 and P63, 46.2 % of the *SOD1*^*G93A*^ mice were unable to follow the 20 cm/s treadmill speed (median = 70; Gehan–Breslow–Wilcoxon test P< 0.0001; n = 11) (Fig. 3B; Video 1 and 2). This locomotor deficit persisted throughout the observation period. This is unlike age-matched control littermates that all were able to follow 20 cm/s treadmill speed (2-way ANOVA and Dunnett’s *post hoc*; P63, P= 0.0060; P70, P= 0.0085; P77, P= 0.0027; P84 P< 0.0001; *wt* n = 8, *SOD1*^*G93A*^ n = 15) (Fig. 3C). Compared with wild-type littermates, *SOD1*^*G93A*^ mice also showed decreased ability to accelerate while coping with the speed of the belt (2-way ANOVA and Dunnett’s *post hoc*; P= 0.0468; *wt* n = 8, *SOD1*^*G93A*^ n = 15) (Fig. 3D), decreased step frequency (2-way ANOVA and Dunnett’s *post hoc;* P= 0.0167; *wt* n = 8, *SOD1*^*G93A*^ n = 15) (Fig. 3F), and a tendency of making shorter steps (Fig 3E) (stride length, 2-way ANOVA and Dunnett’s *post hoc;* P= 0.2157; wt n = 8, *SOD1*^*G93A*^ n = 15). At the ‘*Onset of locomotor phenotype’*, only 50 % of the *SOD1*^*G93A*^ mice could follow a speed of 15 cm/s (*t test* P< 0.0001; n = 11) (Fig. 4A). This reduction in speed was characterized by a distinct locomotor phenotype which included reduced peak acceleration (*t test* P= 0.0118; n = 11) (Fig. 4B), decrease in stride length (*t test* P= 0.0036) (Fig. 4C) and step frequency (*t test* P= 0.0004) (Fig. 4D), and increased drag on the treadmill (Drag counts, *t test* P= 0.0196; Drag duration, *t test* P= 0.0068) (Fig. 4E–F) compared to controls. Characteristically, the locomotor phenotype was not followed by alterations in the left-right hindlimb coordination. Thus, the left-right hindlimb coordination showed out of phase activity with average phase values close to 180 degrees before and after *Onset of locomotor phenotype’* in individual mice (Fig. 3G–I) and for the population of mice (Watson– Williams test, pre-symptomatic vs onset P= 0.1582; n = 15) (Fig. 4G) similar to wild-type mice (Watson–Williams test, *wt* vs *SOD1*^*G93A*^ pre-symptomatic P= 0.1004; *wt* vs *SOD1*^*G93A*^ onset P= 0.3261; n = 15) (Fig. 4G).

**Figure 3.**
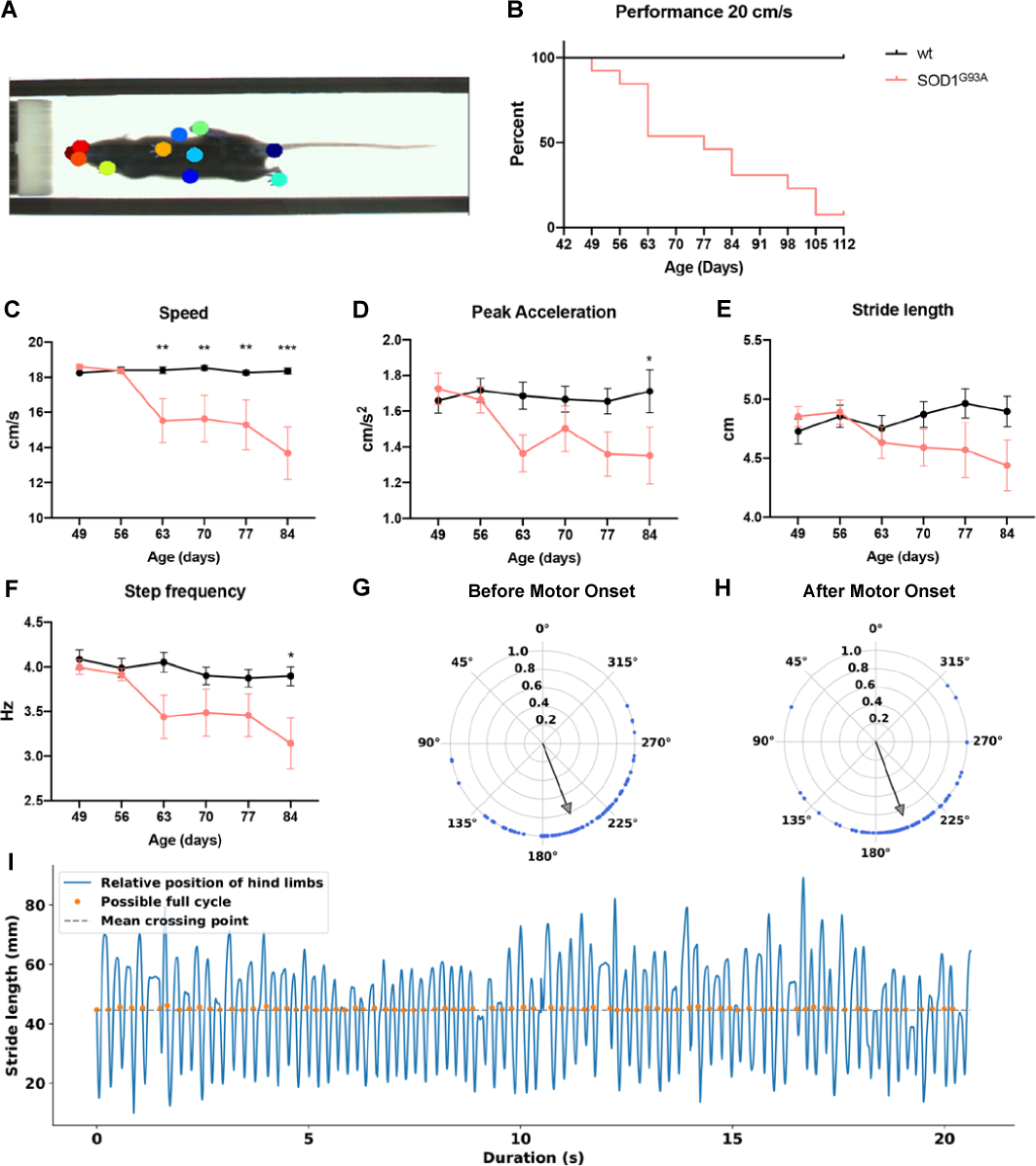
*SOD1*^*G93A*^ mice between postnatal day 49 and 63 show locomotor deficits. (A) Example of tracking approach performed on recorded videos with DeepLabCut analysis tool. Coloured circles show digital markers placed on the animals to extrapolate tracks for analysis. (B) Performance of wild-type (wt) and *SOD1*^*G93A*^ mice on treadmill at a speed of 20 cm/s. Percentage shows that between P49 and P63 46.2 % of the *SOD1*^*G93A*^ mice cannot perform the task (median = 70; Gehan–Breslow–Wilcoxon test P< 0.0001; n = 11). At a longitudinal time scale, mice show a progressive reduction of speed (2-way ANOVA and Dunnett’s *post hoc*; P63, P= 0.0060; P70, P= 0.0085; P77, P= 0.0027; P84 P< 0.0001; wt n = 7, *SOD1*^*G93A*^ n = 15) compared to control mice (C) as well as a progressive reduction in peak acceleration (D) (2-way ANOVA and Dunnett’s *post hoc*; *P= 0.0468; wt n = 7, *SOD-1*^*G93A*^ n = 15), stride length (E) (2-way ANOVA and Dunnett’s *post hoc;* P= 0.2157; wt n = 7, *SOD1*^*G93A*^ n = 15) and step frequency (F)(2-way ANOVA and Dunnett’s *post hoc;* *P= 0.0167; wt n = 7, *SOD1*^*G93A*^ n = 15) when compared to wild-type mice. Hindlimb left-right alternation was compared before (G) and after (H) *“Onset of locomotor phenotype”*. Blue dots represent single steps of a given animal before and after onset. Gray arrows depict mean vectors of average direction of all steps. Phase values at 180 degrees correspond to strict alternation. Stride analysis in (I) shows the out of phase pattern of the hindlimbs during locomotion which was extracted for coordination analysis.

**Figure 4.**
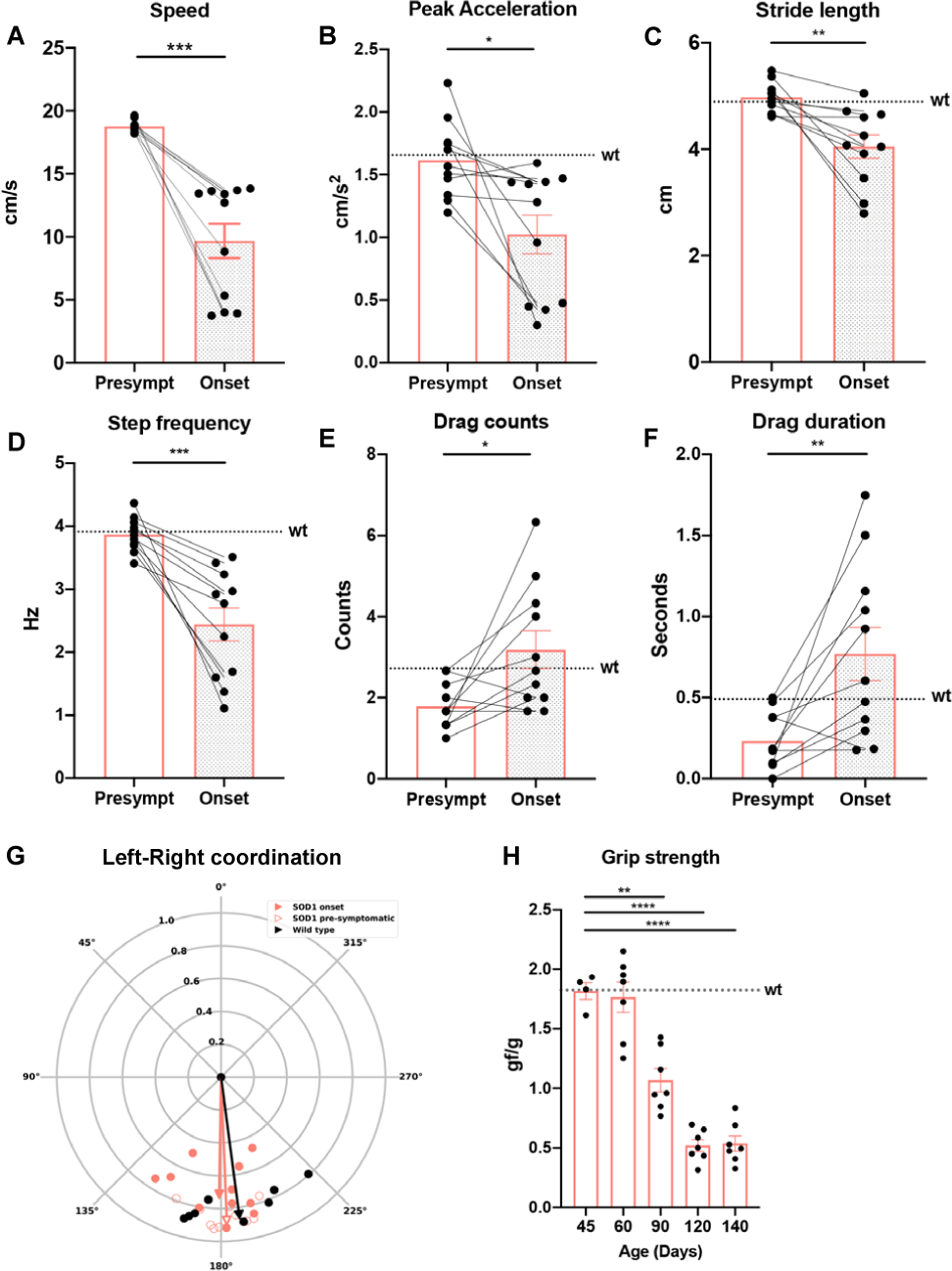
Characterization of the *“Onset of locomotor phenotype”*. Performance of *SOD1*^*G93A*^ mice before and after *“Onset of locomotor phenotype”* is characterized by loss of speed (A) (*t test* P< 0.0001; n = 11), reduction in peak acceleration (B) (*t test* P= 0.0118; n = 11), decrease in stride length (C) (*t test* P= 0.0036; n = 11) and step frequency (D) (*t test* P= 0.0004; n = 11). Moreover, *SOD1*^*G93A*^ mice showed increased dragging events when trying to cope with the speed of the belt, drag counts in (E) (*t test* P= 0.0196; n = 11) and drag duration in (F) (*t test* P= 0.0068; n = 11). Dotted lines show averages for wild-type (wt) mice in all parameters included in the analysis. (G) Quantifications of left-right alternation showed as circular plots. Perfect alternation corresponds to a phase of 180 degrees. Data from individual animals are plotted for each condition (orange-empty = *SOD1*^*G93A*^ pre-symptomatic, orange-full = *SOD1*^*G93A*^ onset, black = wildtype (wt)). The mean vectors for each condition are represented in the respective colors. There is no difference in the mean phase for the different conditions. (Watson–Williams test, pre-symptomatic vs onset P= 0.1582; wt vs *SOD1*^*G93A*^ pre-symptomatic P= 0.1004; wt vs *SOD1*^*G93A*^ onset P= 0.3261; n = 15). (H) Grip strength shows progressive but late manifestation decline of loss low force performance in the *SOD1*^*G93A*^ starting from P90 (one-way ANOVA and Dunnett’s *post hoc*, P90 P< 0.0001; P120 P< 0.0001; P140 P< 0.0001; n = 7 per timepoint).

This progressive *SOD1*^*G93A*^-induced locomotor phenotype, found in half of the animals between postnatal day 49 and 63, paralleled changes in inhibitory inputs to motor neurons but preceded changes at the neuromuscular junctions (NMJs). NMJs in fast twitch fatigable muscles were first affected around P63 (S Fig. 3). The number of fully innervated endplates was significantly reduced at P63 to 58.84 % cof control in the fast tibialis anterior (one-way ANOVA and Dunnett’s *post hoc;* P63, P= 0.0200; n = 4) and 46.94 % in the gastrocnemius muscles (one-way ANOVA and Dunnett’s *post hoc;* P63, P= 0.0003; n = 4) (> 800 NMJs were analyzed per condition at each timepoint – 4 animals per timepoint). In contrast the soleus muscle – composed of 50 % slow-twitch fatigable resistant and 50 % fast-twitch fatigue-resistant fibers – did not show reduction of fully innervated endplates until P84 (one-way ANOVA and Dunnett’s *post hoc;* P< 0.0001; n = 4) (S Fig. 3A–C). Moreover, while the locomotor performance was clearly affected at these early timepoints, low force grip-strength remained unchanged (Fig. 4J). Together these findings show that motor neuron specific affection of motor neuron synapses has measurable changes on motor performance affecting high motor force tasks before low motor force tasks.

### The locomotor phenotype in SOD1^G93A^ mice is mimicked by loss of Engrailed 1 neuron function

Since these changes in locomotor phenotype match the locomotor phenotype previously reported after loss of V1 inter-neuron function in the entire nervous system in wild-type mice observed in isolated spinal cord preparations ^26^, we next tested if V1 interneuron synapse loss on motor neurons is involved in the observed locomotor phenotype in *SOD-1*^*G93A*^ mice. For this we performed experiments with selective dampening of V1 interneurons activity in the spinal cord only. First, we expressed inhibitory (i) DREADDs (hM4Di) – which can be activated by clozapine-*N*-oxide (CNO) leading to hyperpolarization and dampening of cell excitability^41^ – in En1^+^ neurons. To target En1^+^ neurons specifically in the spinal cord, we used a conditional intersectional genetics approach using the *R(ROSA26)C(CAG)::FPDi* mouse line^41^ that allows for dual-recombination with (F; *Flt*) (P; *LoxP*), causing expression of iDREADDs in intersectionally-defined neurons. We therefore crossed *HoxB8*^*FlipO*^ mice^42^ – where *HoxB8* is expressed in the spinal cord from cervical level 4 and downwards – with *En1*^*Cre*^ and *R(ROSA26)C(CAG)::FPDi* mice to obtain triple *HoxB8*^*Flipo*^;*En1*^*Cre*^;*RC::Di* mice (Fig. 5A). Successful intersection of *Cre-Lox* and *Flp-Frt* systems resulted in HA-tag expression in the neuronal population of interest (S Fig. 4). CNO was given intraperitoneally (1mg/kg) as a single dose to triple transgenic mice as well as control mice which were not carrying the *Cre-Lox* and *Flp-Frt* intersectional combination. CNO in this concentration does not have any effect on locomotor performance in wild-type mice^43, 44^. Mice were tested before and up to 20 minutes after administration of CNO on the treadmill with a speed of 20 cm/s (Video 3 and 4). In *HoxB8*^*Flipo*^; *En1*^*Cre*^; *RC::Di* mice CNO effects were detected between 10 and 15 minutes after injection. Seven out of eleven animals responded with a dramatic reduction in speed of locomotion, while the other four showed more pronounced changes in step frequency (Fig. 5; *HoxB8*^*FlipO*^;*En1*^*Cre*^;*RC::Di* n = 11; controls n = 4). Interestingly, the phenotype exhibited by the triple transgenic mice after CNO administration recapitulated the ‘*Onset of locomotor phenotype’* seen in the *SOD1*^*G93A*^ mice. All animals showed reduced speed compared with their performance before injection, and 36.4 % of the animals maintained a speed lower than 15cm/s (one-way ANOVA and Sidak’s *post hoc*, P= 0.0025) (Fig. 5B). Moreover, animals showed a decrease in peak acceleration (one-way ANOVA and Sidak’s *post hoc*, P= 0.0166) (Fig. 5C) reduced stride length (one-way ANOVA and Sidak’s *post hoc*, P= 0.0476) (Fig. 5D) and step frequency (one-way ANOVA and Sidak’s *post hoc*, P= 0.0035) (Fig. 5E). Despite these changes, not all animals dragged on the treadmill to the same extent as *SOD1*^*G93A*^ mice (one-way ANOVA and Sidak’s *post hoc*, Drag counts, P= 0.1168; Drag duration, P= 0.1100) (Fig. 5F–G). Similar to wild-type animals *HoxB8*^*FlipO*^;*En1*^*Cre*^;*RC::Di* left-right hindlimb coordination had phase values around 180 degrees corresponding to alternation and there was no change after CNO administration (Watson–Williams test, P= 0.2876; n = 15 – number of steps per mouse) (Fig. 5H). Interestingly, CNO administration did not change hindlimb grip strength *HoxB8*^*Flipo*^*En1*^*Cre*^ *RC::Di;* grip strength was not significantly different before and after drug administration, and comparable to control animals (one-way ANOVA and Sidak’s *post hoc*, P= 0.6486) (Fig. 5I). Altogether these results show that reducing the activity of spinal En1^+^ neurons leads to a reversible slowing of the locomotor speed in intact mice, strikingly similar to that observed in *SOD1*^*G93A*^ mice.

**Figure 5.**
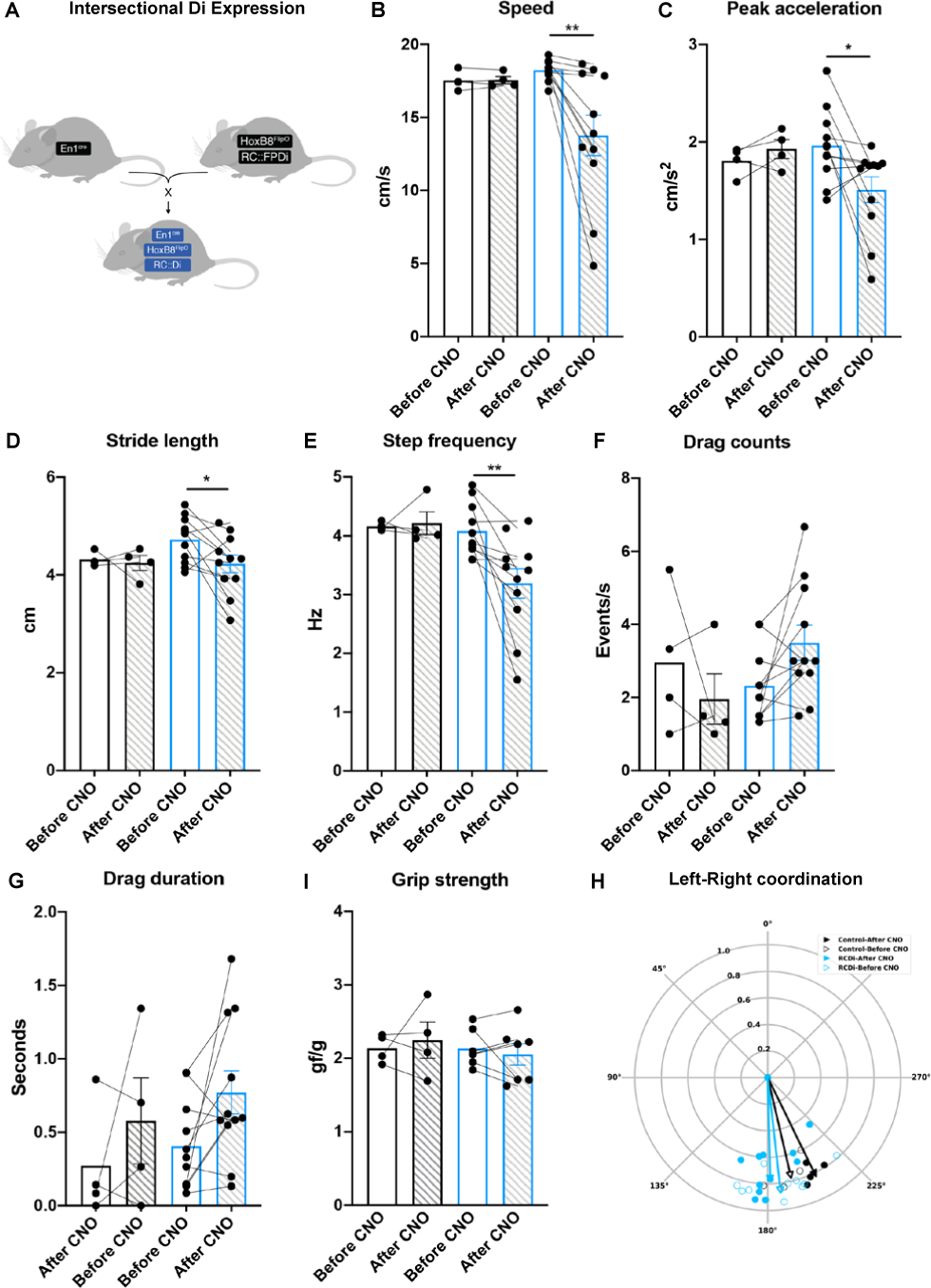
Dampening of spinal V1 interneuron activity recapitulates the *“Onset of locomotor phenotype”*. (A) Mouse genetic approach used to express inhibitory DREADDs specifically in En1^+^ spinal interneurons. *En1*^*cre*^ mouse strain was crossed with *HoxB8*^*FlipO*^;*RC::FPDi* animals. After CNO administration, *En-1*^*cre*^;*HoxB8*^*FlipO*^;*RC::FPDi* mice (blue bars) show loss of locomotor speed (B) compared to their performance before administration of CNO (one-way ANOVA and Sidak’s *post hoc*, P= 0.0025; n = 11). Dual conditional mice also showed decrease of peak acceleration (C) (one-way ANOVA and Sidak’s *post hoc*, P= 0.0166; n = 11) and reduction in stride length (D) (one-way ANOVA and Sidak’s *post hoc*, P= 0.0476; n = 11) and step frequency (E) (one-way ANOVA and Sidak’s *post hoc*, P= 0.0035; n = 11). Despite the loss in speed, not all animals showed increase in dragging events, neither when considering number of drags (F) (one-way ANOVA and Sidak’s *post hoc*, P= 0.1168; n = 11), nor drag duration (G) (one-way ANOVA and Sidak’s *post hoc*, P= 0.1100; n = 11). Grip strength remained unchanged before and after CNO administration (I) (one-way ANOVA and Sidak’s *post hoc*, P= 0.6486; n = 7) as well as left-right alternation (blue-empty = *En1*^*cre*^;*HoxB-8*^*FlipO*^;*RC::Di* before CNO; blue-full = *En1*^*cre*^; *HoxB8*^*FlipO*^;*RC::Di* after CNO; black-empty = control before CNO; black-full = control after CNO) (H) (Watson–Williams test, P= 0.2876; n = 15). CNO administration did not have any effect on control animals (black bars and black arrows; n = 4).

### Dampening of En1^+^ interneuron activity in SOD1^G93A^ mice after ‘Onset of locomotor phenotype’ does not change the locomotor phenotype

We next set out to evaluate the contribution of V1 neurons to the locomotor phenotype in *SOD1*^*G93A*^ mice after their *‘Onset of locomotor phenotype’*. We hypothesized that if the loss of V1 neuron connectivity is the main cause of the locomotor phenotype observed in SOD1^*G93A*^ mice no further worsening of the phenotype should be observed by dampening V1 activity after ‘Onset of locomotor phenotype’ (a phenotypic pathway interaction). To test this, we crossed *HoxB8*^*Flipo*^; *En-1*^*Cre*^; *RC::FPDi* mice with *SOD1*^*G93A*^ mice (Fig. 6A). Of twelve SOD1^*G93A*^ positive mice, four carried both the *Cre-Lox and Flp-Frt* intersectional combination and the remaining littermates were used as control. The *HoxB8*^*Flipo*^; *En1*^*Cre*^; *RC::FPDi; SOD1*^*G93A*^ exhibited the same life span as *SOD1*^*G93A*^ mice (S Fig. 4; 154.34±13.3 days). The *SOD1*^*G93A*^; *HoxB8*^*Flipo*^; *En1*^*Cre*^; *RC::FPDi* and littermate controls were tested weekly on a treadmill from P42 until P98 (S Fig. 4) (median = 77; Gehan– Breslow–Wilcoxon test P< 0.0001; n = 11). During this time all *SOD1*^*G93A*^ animals showed *‘Onset of locomotor phenotype’* which also in this case was characterized by decrease in speed (*t test* P< 0.0001; n = 11) (S Fig. 5A), reduced peak acceleration (*t test* P= 0.0060; N = 11) (S Fig. 5B), decreased stride length (*t test* P= 0.0002), decreased step frequency (*t test* P= 0.0020) as well as increased number of dragging events (*t test;* Drag counts, P= 0.0034; Drag duration, P= 0.0195), and remaining left-right hindlimb alternation not different from controls (Watson–Williams test, pre-symptomatic vs onset P= 0.2792; n = 15 – number of steps) (S Fig. 5). On the day the animals reached the *‘Onset of locomotor phenotype’*, 1mg/kg CNO was administrated as a single dose and mice were tested 10–15 minutes after injection at a speed suitable for their phenotype. *SOD1*^*G93A*^ mice with no iDREADD receptor exhibited no change in any locomotor parameters after CNO administration (Fig 6B–I, orange bars). Similarly, the locomotor phenotype of the *SOD1*^*G93A*^; *HoxB8*^*Flipo*^; *En1*^*Cre*^; *RC::Di* mice did not differ before and after CNO administration; speed (Fig. 6B, magenta bars), stride length (Fig. 6D, magenta bars) and step frequency (Fig. 6E, magenta bars) remained unchanged after administration of CNO. Moreover, the peak acceleration (Fig. 6C, magenta bars), drag events (Fig. 6F–G magenta bars), and left-right hindlimb coordination remained unaffected by CNO administration (Watson–Williams test, P= 0.3682, n = 15 – number of steps per mouse) (Fig. 6I, magenta bars). CNO administration did not cause any changes in grip strength (Fig. 6H). Altogether, these results show that the slowing of locomotion caused by V1 neuron inactivation by inhibitory iDREADDs is occluded in the *SOD1*^*G93A*^ animals after *‘Onset of locomotor phenotype’*, suggesting that the loss of En1^+^ terminals on fast motor neurons is the direct cause of the slowing of locomotor phenotype in *SOD1*^*G93A*^ mice.

**Figure 6.**
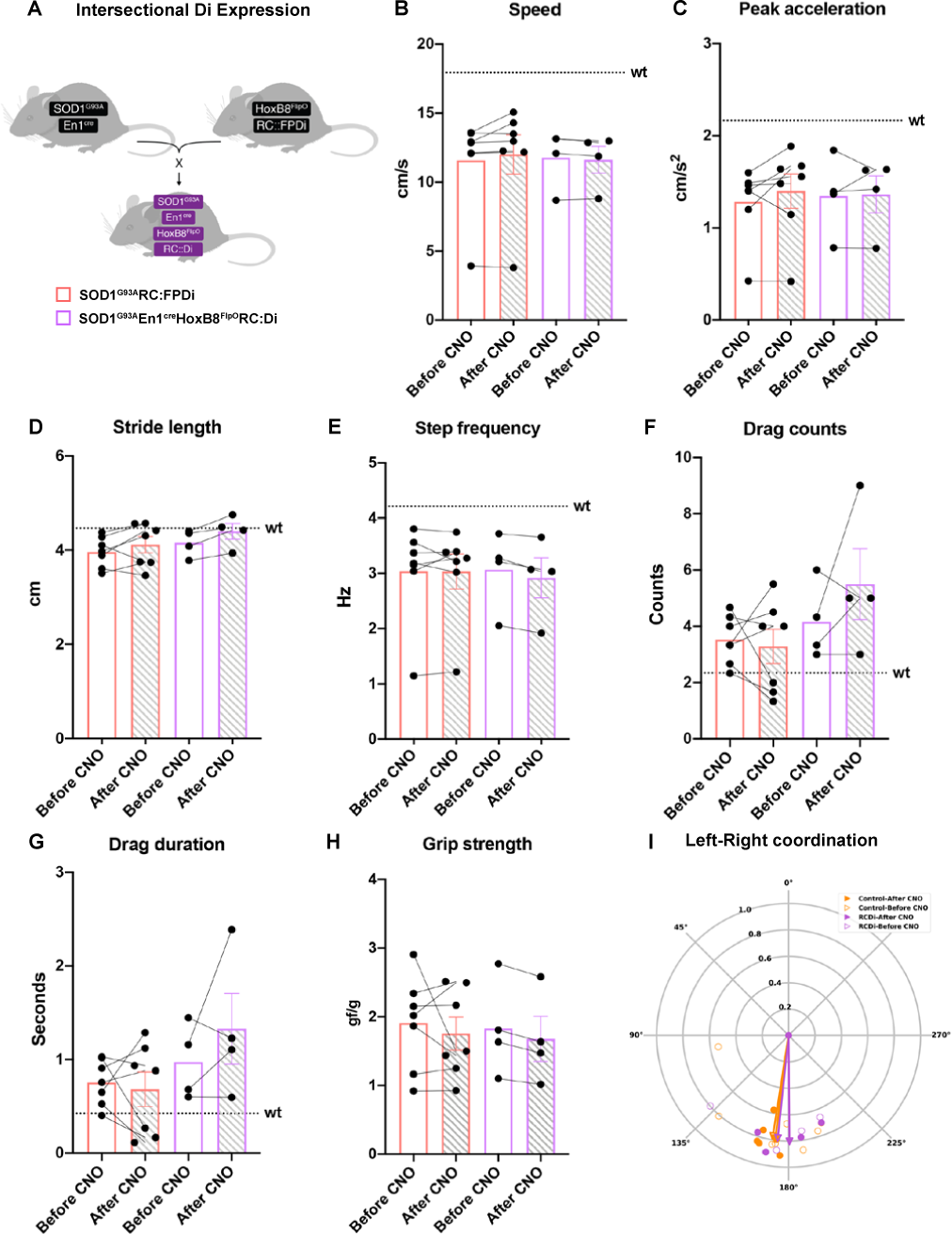
Dampening of spinal V1 interneuron activity does not have an effect in *SOD1*^*G93A*^ mice after *“Onset of locomotor phenotype”*. (A) Mouse crossing utilized to assess specific spinal V1 interneuron silencing in a *SOD1*^*G93A*^ mouse model. *SOD1*^*G93A*^;*En-1*^*cre*^ mice were crossed with *HoxB8*^*FlipO*^;*RC::FPDi. SOD1*^*G93A*^ mice which did not carry the intersectional expression were used as controls. After CNO administration quadruple transgenics (magenta bars) did not show changes in speed (B) (one-way ANOVA and Sidak’s *post hoc*, P= 0.9977; control n = 7; quadruple transgenics n = 4) nor in peak acceleration (C) (one-way ANOVA and Sidak’s *post hoc*, P= 0.9982; control n = 7; quadruple transgenics n = 4). Also stride length (D) (one-way ANOVA and Sidak’s *post hoc*, P= 0.5915; control n= 7; quadruple transgenics n= 4) and step frequency (E) remained unchanged (one-way ANOVA and Sidak’s *post hoc*, P= 0.9602; control n = 7; quadruple transgenics n = 4). *SOD1*^*G93A*^;*En1*^*cre*^;*HoxB8*^*FlipO*^;*RC::Di* mice did not show more dragging events (drag counts, one-way ANOVA and Sidak’s *post hoc*, P= 0.4302; drag duration, one-way ANOVA and Sidak’s *post hoc*, P= 0.5068; control n = 7; quadruple transgenics n = 4). CNO administration did not have any effects on grip strength in *SOD-1*^*G93A*^;*En1*^*cre*^;*HoxB8*^*FlipO*^;*RC::Di* mice (one-way ANOVA and Sidak’s *post hoc*, P= 0.9380; control n = 7; quadruple transgenics n = 4). *SOD1*^*G93A*^ controls not carrying dual intersectional expression did not show any changes after CNO administration (orange bars). Dotted lines show averages for wild-type (wt) animals in all parameters included in the analysis. (I) Left-right alteration remained unaltered after CNO administration in both *SOD1*^*G93A*^;*En1*^*cre*^;*HoxB8*^*FlipO*^;*RC::Di* mice and *SOD1*^*G93A*^ controls (Watson–Williams test, P= 0.3682, n= 15).

## Discussion

The present study uncovers an asymmetric innervation pattern by inhibitory glycinergic spinal interneurons on fast and slow motor neurons. Preferential innervation of fast versus slow motor neurons was selectively lost during early ALS disease stages, both for glycinergic spinal neurons and specific to inhibitory V1 neurons. This motor neuron specific synapse loss mirrors the preferential vulnerability of fast motor neurons in ALS when compared to the slow motor neurons. Remarkably, the loss of inhibitory synaptic inputs is paralleled by a distinct locomotor phenotype in *SOD1*^*G93A*^ mice which is phenocopied by the reversible dampening of V1 interneuron activity in wildtype mice, but it is occluded in *SOD1*^*G93A*^. mice after onset of the locomotor phenotype. These findings reveal a distinct locomotor phenotype in ALS that is linked to V1 neuron synapse retraction on fast motor neurons during early stages of the disease. This retraction appears before motor neuron death and changes in the neuromuscular junctions, suggesting a potential source of the difference in vulnerability of fast and slow motor neurons within the spinal cord in ALS.

Several previous synaptic density investigations performed by immunohistochemistry in *SOD1*^*G93A*^ mice have demonstrated a reduction in glycinergic but not GABAergic inputs onto a-motor neurons starting from postnatal day 60 ^15, 16, 17, 18, 20^. These studies have not, however, differentiated between inputs to fast and slow motor neurons, and therefore were unable to reveal the two folds higher innervation of fast versus slow, and the selective loss of inhibitory synapses onto fast motor neurons. Notably the changes we observed were present also when the synaptic density was normalized to motor neuron area. The motor neuron specific loss is similar for the inhibitory last order V1-population of interneurons, which is composed of 80 % glycinergic neurons and makes up more than 50 % of soma-near motor neuron synapses^27, 35^. These findings suggest that the V1 population of neurons make up a substantial part of the synapse retraction, although further studies are needed to reveal if alterations in connectivity are restricted to the glycinergic V1 interneurons. However, we did not observe alterations in left-right alternation after *‘Onset of locomotor phenotype’* in *SOD1*^*G93A*^ mice, thus, glycinergic commissural V0 interneurons do not appear to be affected at this timepoint. It remains to be investigated if the V1 premotor inputs to slow and fast motor neurons belong to different subpopulations of V1 neurons^45^, which could explain the differential retraction.

The pre-symptomatic loss of soma-near inhibitory synapses is likely to reduce the inhibitory/excitatory synapse balance on fast motor neurons compared to slow motor neurons which may exacerbate the unbalanced excitability in motor neurons observed in pre-symptomatic SOD1^G93A^ mice^46^ and the higher stress responses in vulnerable fast motor neurons in ALS ^3, 5, 7, 47^. Moreover, the recent finding that loss of the Engrailed-1 transcription factor expressed in V1 neurons may lead to motor neuron degeneration^48^ may be another mechanism that contributes to the ALS disease progression when the inhibitory synapses are retracted. The actual cause of the retraction is not known, but our results suggest that ALS may start as an interneuron affection.

A profound finding of this study is that the *SOD1*^*G93A*^ mice exhibited a locomotor phenotype characterized by loss of speed of locomotion and reduction in stride length. This phenotype was observed before motor neuron death and NMJ loss and before loss of grip strength, which suggests that the locomotor phenotype is a sensitive indicator of early ALS progression. These locomotor symptoms have previously been observed in ALS patients with both spinal and bulbar onsets^49^. Notably, the double conditionals *En1*^*Cre*^;*HoxB8*^*FlipO*^;*RC::Di* mice after CNO administration had a similar locomotor phe-notype. While these symptoms seem to be specific for V1 interneurons depletion, the higher dragging events could be detected mainly in *SOD1*^*G93A*^ animals after *‘Onset of locomotor phenotype’* and not in the double conditional mice after CNO administration. This discrepancy suggests that the dragging itself is due to the differences in muscle innervation between *SOD1*^*G93A*^ and wild type mice, since the progressive muscle denervation could affect the capability of the *SOD1*^*G93A*^ mice in sustaining forced locomotion. Notwithstanding, the occlusion of the V1 locomotor phenotype after ‘*Onset of locomotor phenotype’* in ALS strongly suggests that the loss of V1 synapses onto fast motor neurons is the direct cause of the locomotor phenotype observed in ALS since the V1-derived synapses where still present to a large degree on slow motor neurons.

Loss of motor units in tibialis anterior and medial gastrocnemius has been reported from postnatal day 40^4^ and decreased rotarod performance was found from P60^50^. These symptoms have been generally related to changes in muscle innervation, however we found that the V1 synapses withdraw from fast motor neurons before motor neuron degeneration and also before NMJ denervation. Since divergent results were reported on the timepoint of significant NMJ denervation in fast-twitch fatigable muscles by studies utilizing either different (high expressors)^5, 51^ or unspecified^34^ *SOD1*^*G93A*^ mouse models, we analyzed NMJ denervation in tibialis anterior, gastrocnemius and soleus muscles in our *SOD1*^*G93A*^ and *SOD1*^*G93A*^;*GlyT2*^*GFP*^ mice kept on C57Bl6/J congenic background. This analysis demonstrated that glycinergic synapses onto fast motor neurons are lost before NMJ denervation, suggesting that the locomotor phenotype we report is not due to retraction of NMJs but linked to the interneuron changes. Moreover, the grip strength analysis replicated results previously obtained when testing low task forces (significant differences from P90)^50^.

Since the loss of connectivity between V1 inhibitory inter-neurons and fast motor neurons appears during early ALS disease progression in *SOD1*^*G93A*^ mice and leads to locomotor deficits, we suggest that future endeavors could focus on rescuing the synaptic connectivity by stabilizing the V1 soma-near synapses. Thus, stabilization of V1 inhibitory inputs could potentially reduce the stress responses in motor neurons. Several attempts have been directed to stabilize the neuromuscular synapses formed by motor neurons and their target muscles^52, 53, 54, 55^. Drugs known to reduce hyperexcitability directly in motor neurons as the FDA approved Riluzole and Retigabine^56^ have been shown to improve motor neuron survival in ALS. Although, if motor neuron hyper-excitability is also induced by the loss of inhibitory synaptic inputs, attempts to prevent the loss of synaptic inputs early during ALS progression e.g. by overexpressing either genes involved in synaptogenesis or in vesicular trafficking and exocytosis in inhibitory interneurons may be of therapeutic value.

## Supporting information

Video1 SOD1 Pre-symptomatic

Video2 SOD1 Onset

Video3 Before CNO

Video4 After CNO

## Acknowledgements

We thank Dr. Susan Dymecki for providing the conditional *RC::FPDi* mouse. We acknowledge the Core Facility for Integrated Microscopy, Faculty of Health and Medical Sciences, University of Copenhagen. We thank Iryna Vesth-Hansen for technical assistance, and members of Kiehn lab for discussion and comments on previous versions of this manuscript. The authors also thank Peter Simonsson for the drawing used in Figures 2, 5 and 6. This work was supported by the Lundbeck Foundation (IA), the Björklund foundation (IA), the A.P. Møller foundation (IA), the Novo Nordisk Laureate Program (O.K., NNF15OC0014186), The Lundbeck Foundation (OK), the Louise Hansen foundation (RMR), and The Faculty of Health and Medical Sciences (O.K.).

## Authors contribution

Conceptualization IA and OK; Methodology IA, RMR, RS, PL and OK; Investigation: IA and RMR; Writing – Original Draft, IA and OK; Supervision OK; Funding Acquisition, IA and OK.

