## Supplementary material for "Locomotor deficits in ALS mice are paralleled by loss of V1-interneuron-connections onto fast motor neurons"

### Material and methods

*Ethical permits and mouse strains*

All experiments were in accordance with the EU Directive 20110/63/EU and approved by the Danish Animal Inspectorate (Ethical permit: 2018-15-0201-01426). *SOD1*$^{G93A}$ (*B6.Cg-Tg(SOD1-G93A)1Gur/J*) were retrieved from Jackson Laboratory stock no: #004435 and genotyped following the supplier indications, including copy number quantification of the human mutated *SOD1* by qPCR. For qPCRs, the positive control was the *SOD1*$^{G93A}$ founder breeder obtained from Jackson Laboratory carrying 25 copies of the human mutated *SOD1* gene, while a *SOD1*$^{127X}$ strain carrying 19 copies of the human mutated *SOD1* gene[1] was used as negative control. All mouse strains were bred with congenic C57BL6/J animals, stock no: #000664 (Jackson Laboratory). For multiple crossing the following animals kept on C57BL6/J background were used: *GlyT2*$^{GFP}$ [2], *HoxB8*$^{FlipO}$ [3], *R26R-EYFP* stock no: #006148 (Jackson Laboratory). *RC::FPDi* mice[4] were a kind gift from Prof. Susan M Dymecki (Harvard Medical School, Boston, Massachusetts USA). The *En1*$^{cre}$ transgenic mice[5], kept on a C57BL6/J background, were generously obtained from Assistant Prof. Jay Bikoff (St. Jude Children's hospital, St Louis, Texas USA). After multiple crossing, animals' genotype, including hSOD1 copy number, and phenotype were analyzed, including their survival. Mice were housed according to standard conditions: fed *ad libitum*, constant access to water, and a 12:12 hour light/dark cycle. Both males and females were included in the study and non-transgenic *SOD1*$^{G93A}$ littermates were used as controls.

*Intraspinal injections and viral delivery*

For viral transfection of the En1-expressing spinal interneurons, P42 mice were anaesthetized with 2% isoflurane and the lumbar level of the spinal cord was exposed for stereotaxic injections (Neurostar). A small incision was performed with micro-scissors between the T12 and the T13 vertebrae in order to deliver the virus in the lumbar spinal cord. For visualization, virus was mixed with 4% fast green (Invitrogen) dissolved in saline and injected using a glass micropipette at a rate of 100 nl/minute. The micropipette was kept in place for 2 minutes after viral delivery to avoid backflow. Bilateral injections of an AAV1-phSyn1(S)-FLEX-tdTomato-T2A-SypEGFP-WPRE ($5.56 \times 10^{11}$ /ml, Viral Vector Core, Salk Institute for Biological Sciences; Addgene 51509) were performed in *SOD1*$^{G93A}$*;En1*$^{cre}$ mice and *wt;En1*$^{cre}$ littermates to assess synaptic connectivity of En1 positive neurons to fast fatigable and slow motor neurons. Animals were treated post-operatively with Buprenorphine (Tamgesic) diluted in saline at a concentration of 0.3 mg/ml for pain relief. Three weeks after injections, at postnatal day P63, mice were perfused and the tissue was processed for immunohistochemistry (see *spinal cord immunohistochemistry).*

*Spinal cord immunohistochemistry*

Animals were anesthetized with an overdose of Pentobarbital (250 mg/kg) and perfused transcardially with pre-chilled phosphate buffered saline (PBS, Gibco) followed by pre-chilled 4% paraformaldehyde (PFA, HistoLab). Spinal cords were dissected and cryoprotected in 30% sucrose dissolved in PBS. Tissue was then sectioned at 30 μm thickness on a cryostat (Thermo Fisher Scientific). Coronal sections were collected on Superfrost Plus slides (Thermo Fisher Scientific) and then stained by immunohistochemistry. Slides were washed for 10 minutes in PBS at room temperature and blocked for 1 hour in PBS with 0.1% Triton-X100 (PBS-T, Sigma Aldrich) and 1.5% donkey or goat serum (Invitrogen). When utilizing monoclonal mouse antibodies, blocking was performed over night at 4 °C with Fab Fragment anti-mouse (115-007-003, Jackson Laboratory) at a concentration of 1:50 in PBS-T and 1.5% donkey serum to reduce unspecific background. Sections were incubated for 48 hours at 4 °C in primary antibodies diluted in blocking solution (Table 1). Slides were washed three times in PBS at room temperature for 10 minutes each and then incubated for 1 hour with appropriate secondary antibodies (1:500, Alexa Fluor 488, 568, 647, Invitrogen) diluted in blocking solution. Counterstaining was performed either with Hoechst 33342 (1:2000 Invitrogen) or NeuroTrace 435 (1:500, Invitrogen). For synaptic density counterstaining, a Synaptophysin antibody, Alexa Fluor 594 conjugated, was incubated overnight (1:100, sc-17750 Santa Cruz). After three washes is PBS, slides were dried overnight, and cover slipped using Mowiol 4-88 mounting media (Sigma Aldrich). Microphotographs were obtained utilizing either Zeiss LSM 700 or 780 confocal microscopes.

*Image analysis*

All image analysis and quantifications were conducted utilizing Fiji software. For synaptic density quantifications, microphotographs, obtained with a 20x objective – zoom = 1, were transformed to a grayscale 8-bit images and quantified after applying a threshold for signal intensity/background correction. Images with higher background/noise ratio were excluded. ROIs were drawn around the motor neurons of interest that showed clear soma staining and visible nucleus, and the pixel contained in the area of interest were quantified with the Fiji function "Analyze particles". At 20x magnification – zoom=1, synaptic terminals had a size between 1 and 5 pixels, thus a threshold of 5 pixel was used to identify and quantify individual synapses. Masks of all quantified images were created to assure careful inclusion of pixels (Figure 1). Number of synaptic terminals was then normalized for motor neuron areas and averaged for the number of motor neurons quantified in each mouse. When comparing wild-type and *SOD1*$^{G93A}$ littermates the average synaptic density was quantified across control mice and used to normalize the synaptic density in the *SOD1*$^{G93A}$ mice. Between 8-10 images per mouse per condition were analyzed. For neuromuscular junction (NMJ) analysis, z-stack confocal images were acquired, and the maximal projection was used for quantification. Microphotographs were acquired with 20x objective – zoom = 1 and 16 pictures per muscle per mouse per condition were quantified. Depending on the level of innervation, NMJs were categorized as full, partial or empty (Fig. S3).

*NMJ quantification*

For muscle analysis and NMJ quantification, mice were anesthetized by Pentobarbital overdose (250 mg/kg) and sacrificed by decapitation at three different time points: postnatal day 45, 63 and 84. Gastrocnemius, tibialis anterior and soleus muscles were dissected and fixed in 4% PFA for 20 minutes, washed twice in PBS for 10 minutes, and cryoprotected in 30% sucrose. The muscles were then sectioned in a cryostat to obtain 100 μm thick slices which were collected in a 24-well plate (Falcon) for immunohistochemistry. Free floating sections were blocked for 1 hour in PBS-T and incubated for 48 hours at 4 °C in primary antibodies diluted in blocking solution (Table 1). Sections were then washed for 1 hour in PBS-T and incubated for 3 hours in secondary antibody (1:500, Alexa Fluor 555, Invitrogen) at room temperature. After three washes in PBS for 10 minutes at room temperature, sections were incubated with an α-Bungarotoxin Alexa Fluor 488 conjugated antibody (1:500, Invitrogen) for 10 minutes at room temperature. After two 5 minute washes in PBS, sections were mounted on SuperFrost Plus slides, dried overnight and cover-slipped using Mowiol 4-88 mounting media. Microphotographs were obtained utilizing Zeiss LSM 700 confocal microscope.

*DigiGait treadmill test*

Locomotor performance was assessed using the DigiGait motorized transparent treadmill (Mouse Specifics, Inc.), which allows recording of animals from a ventral view. Mice were pre-trained on the treadmill at speeds of 15 and 20 cm/s at postnatal day 42 and then tested weekly from postnatal day 49 to 112 for analysis of disease progression. For inhibitory chemogenetic experiments (activation of inhibitory DREADD receptors), age matching animals were tested before and after administration of clozapine-N-oxide (CNO). After a 2 minute acclimatization to the treadmill, mice were recorded at a speed of 20 cm/s for 10 seconds in 3 consecutive trials with 2 minute rest periods in between recordings. Belt speed was adjusted to 15, 10 or 5 cm/s as needed, depending on the locomotor capability.

*Digital Tracking Analysis*

The videos captured during the treadmill experiments were analyzed in DeepLabCut (DLC)[6] to extract tracks based on digital markers placed on the animals (Fig.3-A). Eleven digital markers were placed on the mice in a total of 50 frames. These labelled frames extracted from a subset of the videos created the training labels used for automatic tracking in DLC. A pre-trained ResNet-50 (https://arxiv.org/abs/1512.03385) model provided in DLC was further fine-tuned with these labels and trained for a total of 4500 iterations. This fine-tuned model was then used to predict tracks from all the videos.

The tracks output by DLC were further analyzed to extract the locomotor measures reported in the manuscript. Details of estimating the reported measures are described below.

We extracted the horizontal displacement of the animals $x_n$ on the treadmill, collected them into a vector denoted $X^i = [x_1, \ldots, x_N]$ where $n = [1, .., N]$ and $N$ is the total number of frames in the video. Individual mice are indexed with superscript $i$, which is dropped in the remaining description for ease of notation.

**Speed:** Position estimates from all the markers, except the four used to track the paws, were utilised to estimate instantaneous speed of the animals. Difference in position, $\Delta x_n$, between the tracked markers in successive frames and the inter-frame duration, $\Delta T$, were used to estimate the instantaneous speed per tracked marker:

$$v_m = \frac{\Delta x_n}{\Delta T} \, (cm/s) \text{ where } m = [1, \ldots, 7].$$

The estimates from the seven markers were used to obtain the average instantaneous speed at time $t$ in the video, $v_t$:

$$v_t = \frac{1}{7} \sum_m v_m \, (cm/s).$$

A moving average filter of window size 10 was used to reduce high frequency noise in the speed estimates. The average speed per video was estimated by computing the mean over the instantaneous speed throughout the video:

$$v = \frac{1}{N} \sum_t v_t \, (cm/s).$$

**Acceleration:** Sudden increase or decrease in the speed of the animals were captured by estimating the instantaneous acceleration derived from the instantaneous speed estimates:

$$a_t = \frac{\Delta v_t}{\Delta T} \, (cm/s^2)$$

We set a threshold of $0.25s$ of continuous deceleration to detect the event as a *drag* and conversely a duration of $0.25s$ of continuous acceleration as a *recovery* event. The maximum acceleration attained by the mice in each video is reported as *peak acceleration*:

$$a_P = max(a_t) (cm/s^2)$$

These acceleration measures are reported in Figures 4-B, 4-E, 4-F.

**Coordination:** The tracks for the four paws were used to estimate the *stride length* and *step frequency* of the mice. We first estimated the total number of steps, $S$, in the video by analysing the relative position between the left and the right hind limbs (Figure 3-I). This provided an estimate of the step frequency:

$$F = \frac{S}{T} (Hz)$$

where $T$ is the video duration.

Stride length, $L$ was estimated from the average speed and step frequency:

$$L = vF \, (cm).$$

For each step, the phase difference between the left and right limbs, $\Phi_s$ where $s = [1, \ldots, S]$, were computed from the cadence plot in (Figure 3-I). The individual phase values are then plotted on a circle representing the interval of possible phases from 0 to 360 degrees. The average phase difference over all the steps per animal, $\Phi$, were computed using circular statistics[7]. The average phase is indicated by the direction of the vector originating from the center of the circle. Phase values around 180 degrees indicate alternation while phase values around 0 (or 360) degrees indicate synchrony. The individual animal level coordination profiles were further aggregated over all animals within a study group to report the group level coordination as seen in all the circular plots (Figures 4-G, 5-H, 6-I).

*Grip strength test*

Low force motor performance was assessed by measuring hind limb grip strength. Measurements were performed as previously described[8] using a computerized grip strength meter (Cat. No. 47200, Ugo Basile). *SOD1*$^{G93A}$ animals were tested at postnatal days 45, 60, 90, 120 and 140 for analysis of disease progression, as well as before and 10-15 minutes after administration of CNO for analysis of chemogenetic inhibition of En1 spinal interneurons. The peak force was recorded in gram-force (gf) and normalized to body mass. All measurements were performed in triplicates.

*Chemogenetic experiments*

For experiments utilizing inhibitory DREADDs, both males and females *En1*$^{cre}$*;HoxB8*$^{FlipO}$*;RC::FPDi* mice between postnatal day 63 and 90 were used. Animals were previously trained in the treadmill at 15 and 20 cm/s speeds and then tested as described above before and after injection. Also for grip strength test, animals were previously trained and then tested before and after CNO delivery. Clozapine-N-oxide (CNO, Tocris, #4936) was diluted in saline at a concentration of 1 mg/kg and delivered intraperitoneally. Animals were then tested every 5 minutes after injections up to 20-30 minutes.

*Statistics*

All statistical analysis was performed with GraphPad software unless otherwise specified. No statistical methods were used to pre-determine sample sizes; our sample sizes are similar to those reported in previous publications[9, 10, 11]. Mice were randomly allocated to different groups for the *in vivo* experiments using a block design. Data collection was performed blind to the conditions of the experiments although, in behavioral assessment, ALS mice could be easily recognized at later stages. All tested animals were included in the study, except for one *SOD1*$^{G93A}$ excluded from the survival assessment due to the appearance of wounds that required sacrifice before humane end-stage was reached. Unpaired Student's t test was used to compare two groups and one-way ANOVA was used for multiple comparisons. When taking into account multiple variables, two-way ANOVA was used. ANOVA tests were followed by appropriate post hoc analysis as indicated in the figure legends. All results are expressed as mean±SEM, reported n values represent distinct biological replicates. $P < 0.05$ was considered statistically significant, asterisks in figures and figure legends are * $P < 0.05$, ** $P < 0.01$ or *** $P < 0.001$. To compare the left-right coordination during disease progression and before and after treatment with CNO we used Watson-Williams test using 15 steps per mouse.

*Data availability*

The data that support the findings of this study are available from the corresponding authors upon reasonable request.

*Code availability*

The code used to analyze data and produce figure content is available from the corresponding authors upon request.

*Table 1. Antibody list*

| Application | Target | Source | Host species | Concentration | Reference |
|---|---|---|---|---|---|
| NMJ quantification | α-Bungarotixin Alexa Fluor 488 | Invitrogen B-13422 | | 1:500 | [9, 12] |
| NMJ quantification | Neurofilament (165 kDa) | DSHB (2H3) | Mouse | 1:50 | [9, 12] |
| NMJ quantification | SV2A | DSHB (SV2) | Mouse | 1:50 | [9, 12] |
| Synaptic density | GFP | Abcam Ab-13970 | Chicken | 1:1000 | [13] |
| Synaptic density | MMP-9 | Sigma M9570-100UG | Goat | 1:1000 | [14] |
| Synaptic density | ErrBeta | R&D PP-H6705-00 | Mouse | 1:500 | [15] |
| Synaptic density | Synaptophysis Alexa Fluor 594 | Santa Cruz Sc--17750 | Mouse | 1:100 | [16] |
| Synaptic density | DsRed | Clontech 632496 | Rabbit | 1:1000 | [17] |
| Intersectional expression | HA-tag | Sigma Aldrich H6908 | Rabbit | 1:100 | [18] |
| Intersectional expression | NeuN | Millipore ABN91 | Chicken | 1:1000 | [19] |

All primary antibodies have been extensively used and evaluated for specificity in many previous publications as indicated in the Antibody Registry - https://antibodyregistry.org and from the manufactories (Abcam, Sigma, Clontech, Millipore, Invitrogen, DSHB, Santa Cruz)) homepages. Here references are given to a few studies.

*References*

10. Cregg JM, Leiras R, Montalant A, Wanken P, Wickersham IR, Kiehn O. Brainstem neurons that command mammalian locomotor asymmetries. *Nat Neurosci*, (2020).

**Supplementary Figure Legend**

**Supplementary figure 1. Characterization of *$SOD1^{G93A}$;$GlyT2^{GFP}$* animals**. (A) Copy number of human SOD1 mutations carried by the animals included in the study. Orange dots depict fold change for *$SOD1^{G93A}$* mice, green dots for the *$SOD1^{G93A}$;$GlyT2^{GFP}$* mice. First orange dot is the positive control - *$SOD1^{G93A}$* founder carrying 25 copies of the mutated gene, second gray dot the negative control - S*$OD1_{127X}$* carrying 19 copies of the mutated gene. (B) Weights of *$SOD1^{G93A}$* and *$SOD1^{G93A}$;$GlyT2^{GFP}$* males compared to wild-types. After crossing, *$SOD1^{G93A}$;$GlyT2^{GFP}$* mice do not differ from the *$SOD1^{G93A}$* ones but both strains differ from wild-type littermates (Multiple t test, P112 P=0.0459; n=8). Motor neuron quantification was performed utilizing Nissl staining in P84 wild-type (C) and *$SOD1^{G93A}$;$GlyT2^{GFP}$* (D) mice. At this time point changes in motor neuron survival could not be detected (D) (*t test,* P=0.2278; n=3) as well as motor neuron shrinkage (E) (*t test,* P=0.1980; n=3). Fluor Nissl in cyan, scale bar in (C) = 100 μm.

**Supplementary figure 2. Characterization of *$SOD1^{G93A}$;$En1^{cre}$* animals.** (A) Spinal cord cross section from *$SOD1^{G93A}$;$En1^{cre}$;$Rosa26^{YFP}$* mice at P63. Scale bar = 50 μm. Despite somata were clearly $YFP^+$, synaptic terminals could not be visualized in this reporter mouse line (B) MMP9 in blue, synaptophysin (SYP) in red, YFP in green. (C) Copy number quantifications of the human mutated *SOD1* gene carried by *$SOD1^{G93A}$;$En1^{cre}$* mice used in the study. Orange dots = *$SOD1^{G93A}$*, purple dots = *$SOD1^{G93A}$;$En1^{cre}$* and black dot = negative control S*$OD1^{127X}$*. (D) Weights of *$SOD1^{G93A}$, $SOD1^{G93A}$;$En1^{cre}$* males compared to wild-type mice. *$SOD1^{G93A}$* and *$SOD1^{G93A}$;$En1^{cre}$* mice show similar progression in weight loss over time when compared to their wild-type littermates (Multiple t test, P120 P=0.0117; n=8).

**Supplementary figure 3. Neuromuscular junction (NMJ) innervation assessment in the fast-twitch fatigable *Tibialis Anterior* (A) and *Gastrocnemius* (B) muscle fibers and in the slow-twitch *Soleus* (C) muscle fibers.** NMJ were categorized either as fully innervated, shown by + in (D), or as partially innervated, shown as * in (D), or as empty, shown as # in (D). Significant denervation in TA (A) (one-way ANOVA and Dunnett's *post hoc,* P63 P=0.0200; P84 P=0.0005; n=4 per condition per time-point) and GN (B) was in P63 mice (one-way ANOVA and Dunnett's *post hoc,* P63 P=0.0003; P84 P=0.0001; n=4 per condition per timepoint). (C) In SOL, significant denervation was much later starting from P84 (one-way ANOVA and Dunnett's *post hoc,* P84

P<0.0001). (E) Weights of the *SOD1$^{G93A}$* mice included in the grip strength analysis, used to normalize force.

**Supplementary figure 4. Characterization of *SOD1$^{G93A}$;En1$^{cre}$;HoxB8$^{FlipO}$;RC::Di mice*.** (A) Dual recombinase of (F; *Flt*) (P; *LoxP*) results in HA-tag expression in *En1$^{cre}$;HoxB8$^{FlipO}$;RC::Di* positive neurons, higher magnification in (B). HA-tag in green, NeuN in red and Hoechst in blue. Scale bar = 100 μm. (C) Weights of *SOD1$^{G93A}$;En1$^{cre}$;HoxB8$^{FlipO}$;RC::Di and SOD1$^{G93A}$* mice do not differ when compared to wild-type mice (Multiple t test, P84 P=0.4458; n=7). (D) Copy number quantifications of the human mutated SOD1 gene carried *by SOD1$^{G93A}$;En1$^{cre}$;HoxB8$^{FlipO}$;RC::FPDi* included in the study. Orange dots (positive control) = *SOD1$^{G93A}$*, purple dots = *SOD1$^{G93A}$;En1$^{cre}$;HoxB8$^{FlipO}$;RC::FPDi* and black dots = negative control *SOD1$^{127X}$*. (E) Phenotype validation of the of the *SOD1$^{G93A}$;En1$^{cre}$;HoxB8$^{FlipO}$;RC::FPDi* showed similar phenotype as *SOD1$^{G93A}$* mice on the treadmill (median=77; Gehan-Breslow-Wilcoxon test P<0.0001; n=11). (F) The locomotor phenotype included progressive loss of speed (two-way ANOVA and Dunnett's *post hoc*, P70 P=0.0013; P77 P=0.0009; P84 P<0.0001; wild-type n=9 and *SOD1$^{G93A}$;En1$^{cre}$;HoxB8$^{FlipO}$;RC::FPDi* n=11), reduced peak acceleration (G) (two-way ANOVA P=0.0013; wild-type n=9 and *SOD1$^{G93A}$;En1$^{cre}$;HoxB8$^{FlipO}$;RC::FPDi* n=11), decrease in stride length (H) (two-way ANOVA, P=0.0094; wild-type n=9 and *SOD1$^{G93A}$;En1$^{cre}$;HoxB8$^{FlipO}$;RC::FPDi* n=11) and in step frequency (I) (two-way ANOVA, P=0.0008; wild-type n=9 and *SOD1$^{G93A}$;En1$^{cre}$;HoxB8$^{FlipO}$;RC::FPDi* n=11).

**Supplementary figure 5. Characterization of *"Onset of locomotor phenotype"* in *SOD1$^{G93A}$;En1$^{cre}$;HoxB8$^{FlipO}$;RC::Di* mice.** As for the *SOD1$^{G93A}$* mice, the *"Onset of locomotor phenotype"* in *SOD1$^{G93A}$;En1$^{cre}$;HoxB8$^{FlipO}$;RC::Di* mice was characterized by decrease in speed of locomotion (A) (*t test* P<0.0001; n=11), reduced peak acceleration (B) (*t test* P=0.0060; N=11), decreased stride length (C) (*t test* P=0.0002; n=11) and decreased step frequency (D) (*t test* P=0.0020; n=11). Dragging events were increased both in number (E) (*t test,* P= 0.0034; n=11) and duration (F) (*t test*, P= 0.0195; n=11). (G) Left-right coordination remained unchanged (Watson-Williams test, pre-symptomatic vs onset P=0.2792; n=15). Magenta-full = onset; magenta empty = pre-symptomatic; black dots = wild-type. Dotted lines show averages for wild-type (wt) animals in all parameters included in the analysis.

**Video 1.** Locomotor analysis of a *SOD1*$^{G93A}$ mouse at pre-symptomatic stage (postnatal day 49). Mouse was placed on a treadmill at a speed of 20 cm/s and filmed at 150 frames per second. Coloured circles in the video indicate digital markers used to extrapolate tracking information of areas of interest of the mouse (snout, torso, paws, tail). Visualization of real time analysis shows profiles for speed, acceleration and cadence. Circular plot depicts phases of left-right alternation (perfect alternation 180 degree), mean vector changes depending on the phase of the consecutive steps. Video is shown at 40 frames/s.

**Video 2.** Analysis of the same *SOD1*$^{G93A}$ mouse showed in Video 1 after *"Onset of locomotor phenotype"* (postnatal day 63). Due to locomotor deficits the speed was reduced to 10 cm/s. Mouse was placed on a treadmill and filmed at 150 frames/s. Real time analysis of digital markers shows differences in speed, acceleration and cadence profiles; circular plot shows phases of left-right alternation during dragging events. Video is shown at 40 frames/s.

**Video 3.** Locomotor analysis of an age-matching *En1*$^{cre}$*;HoxB8*$^{FlipO}$*;RC::Di* mouse before CNO administration. Panels represent speed, acceleration and cadence profiles of the mouse placed on a treadmill at a speed of 20 cm/s. Video was recorded at 150 frames/s. Phases of left-right alternation are depicted on the circular plot, mean vector is represented by the arrow. Video is shown at 40 frames/s.

**Video 4.** Analysis of the same *En1*$^{cre}$*;HoxB8*$^{FlipO}$*;RC::Di* mouse showed in Video 3 after administration of Clozapine-N-Oxide (1 mg/kg) i.p. Fifteen minutes after silencing of spinal En1 interneurons the mouse can walk on a treadmill only at a speed of 15 cm/s. Panels show changes in speed, acceleration and cadence. Left-right alternation depicted in circular plots remains unchanged after CNO administration. Video was recorded at 150 frames/s and is shown at 40 frames/s.

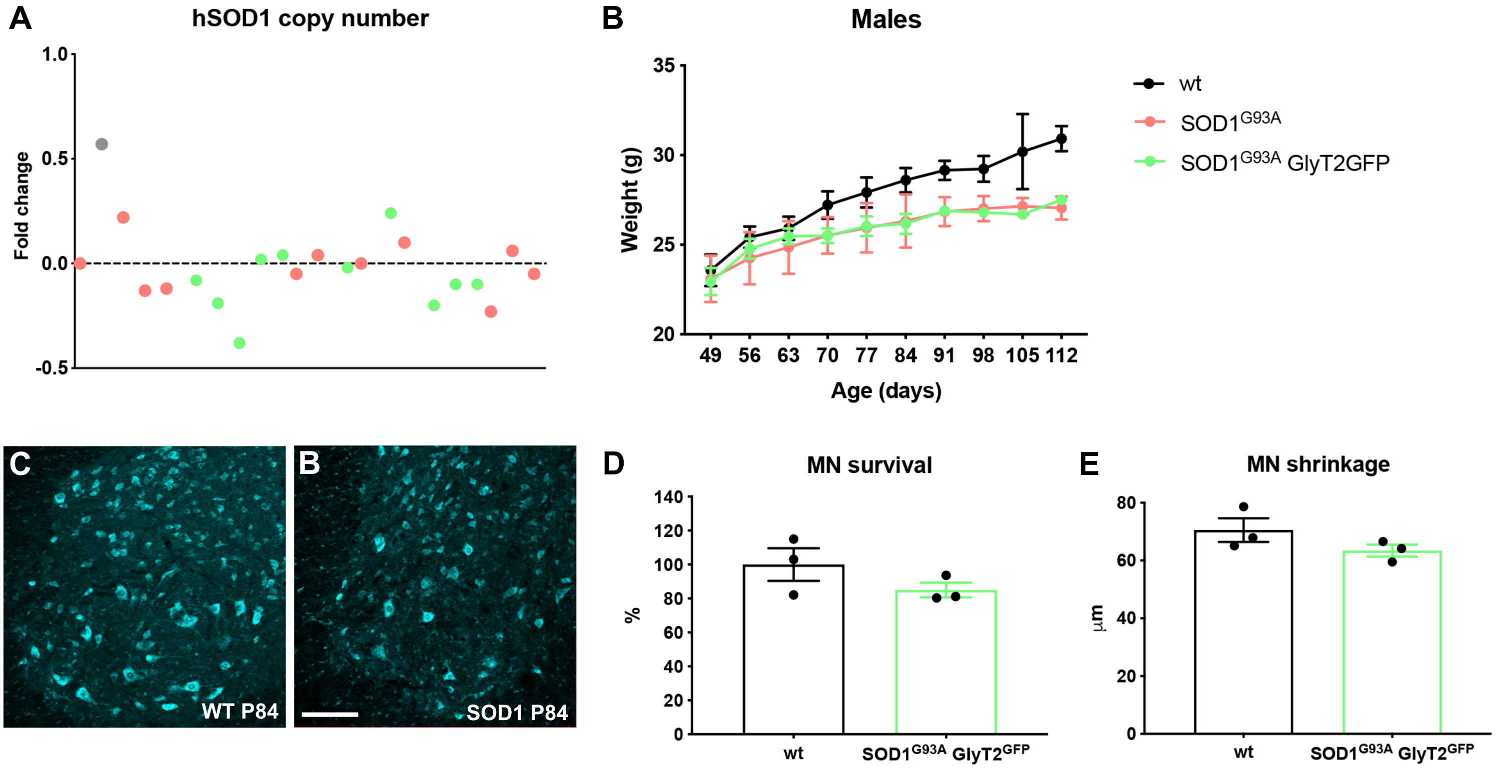


Supplementary Figure 1

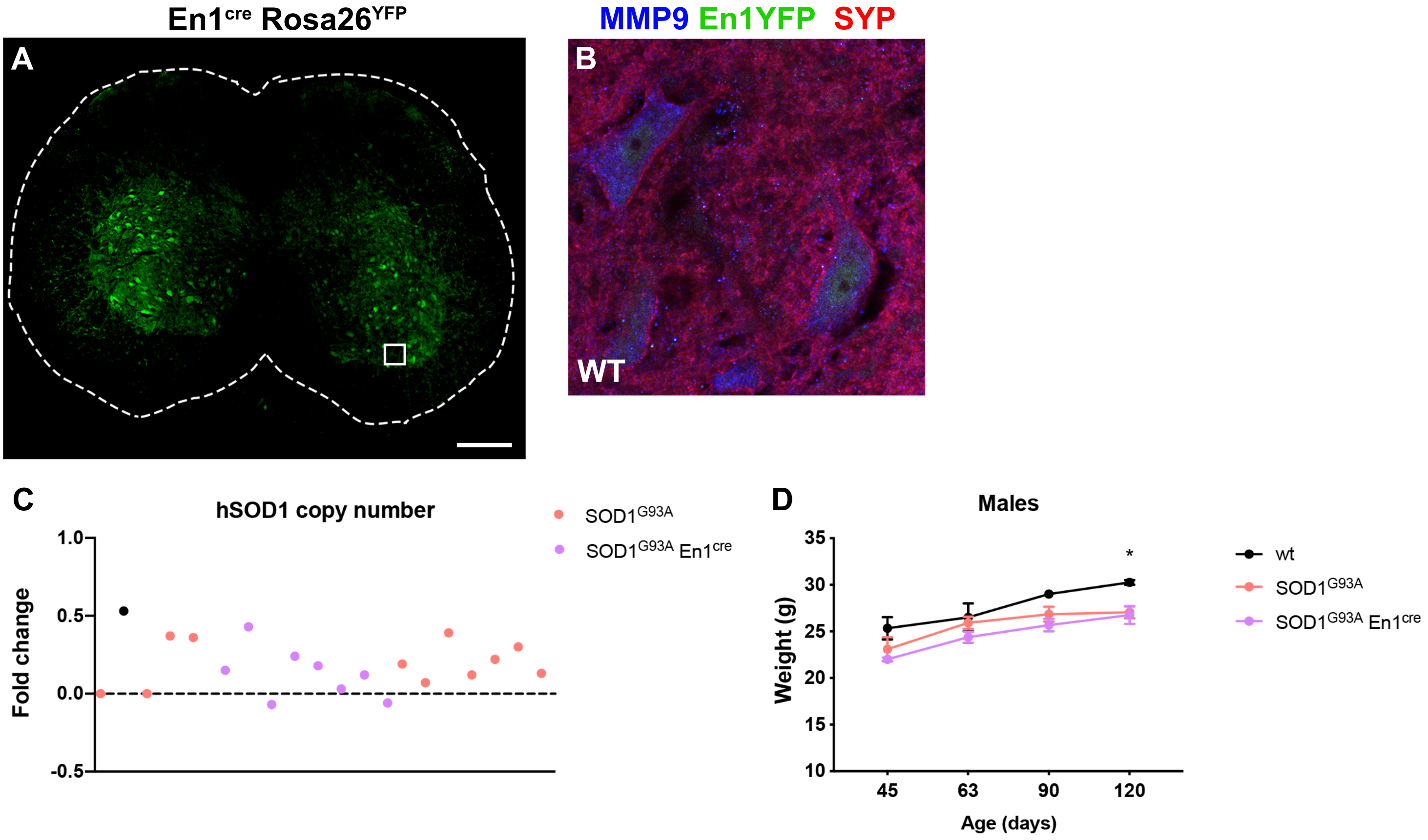


**Supplementary figure 2**

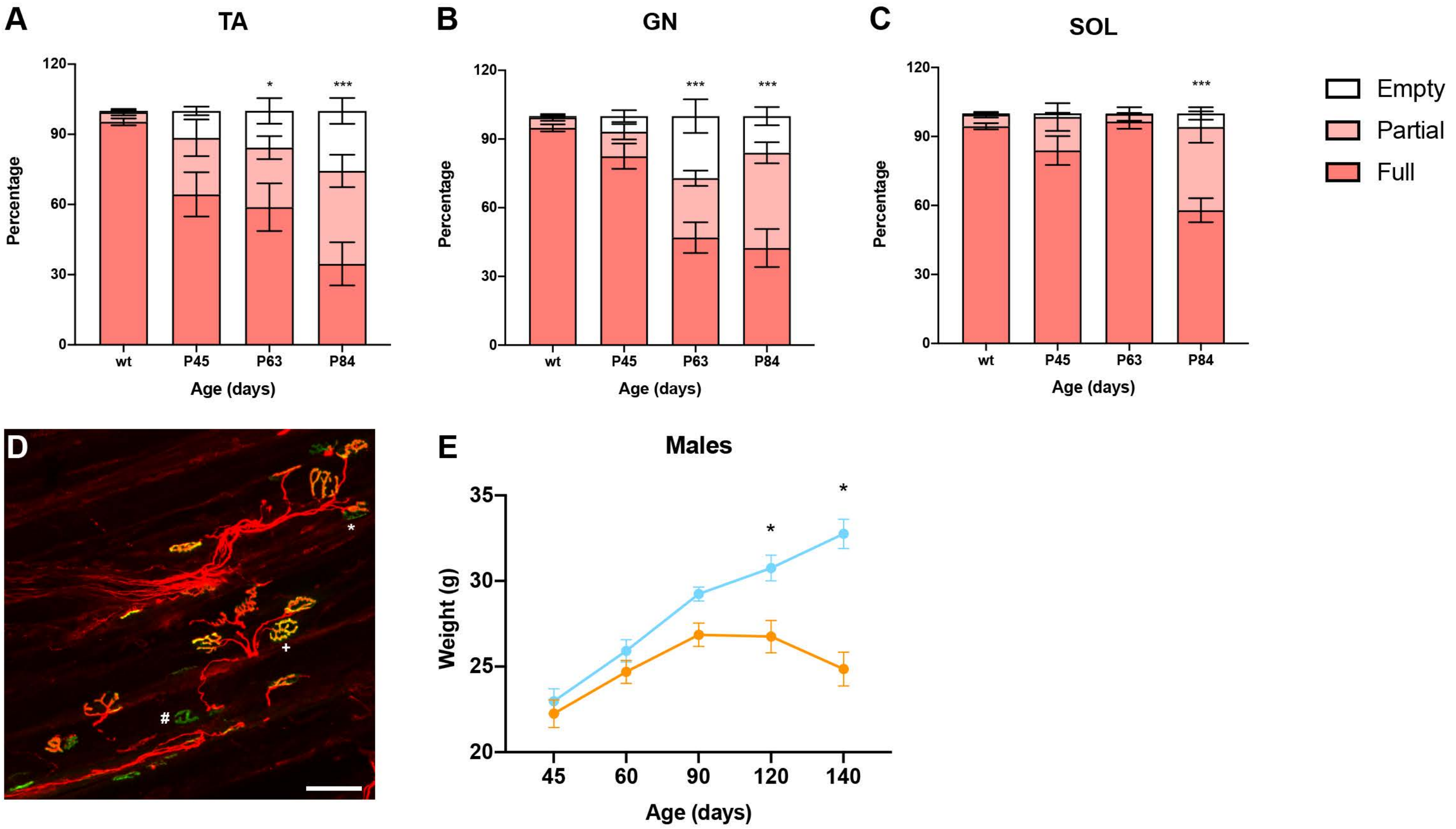


**Supplementary figure 3**

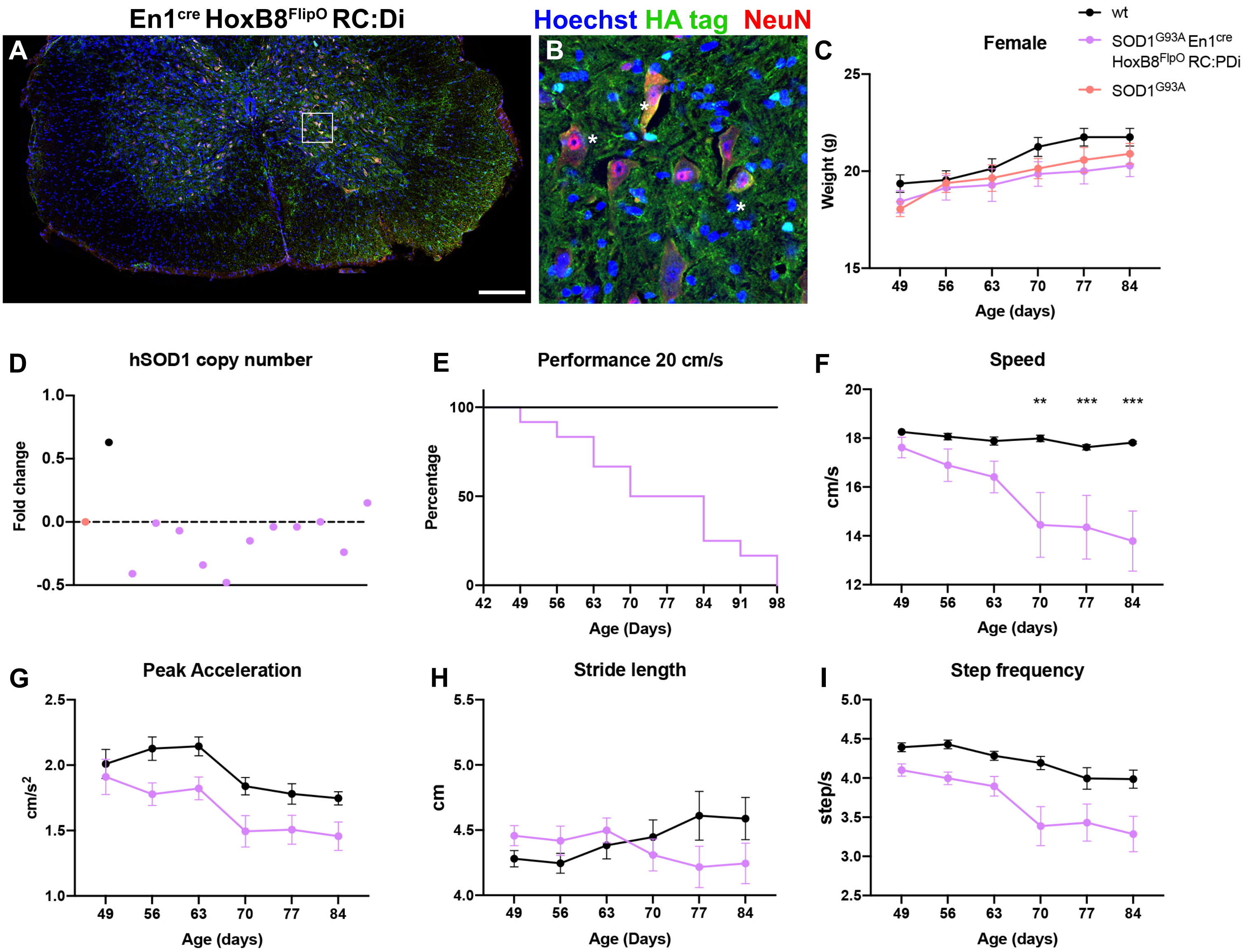


**Supplementary figure 4**

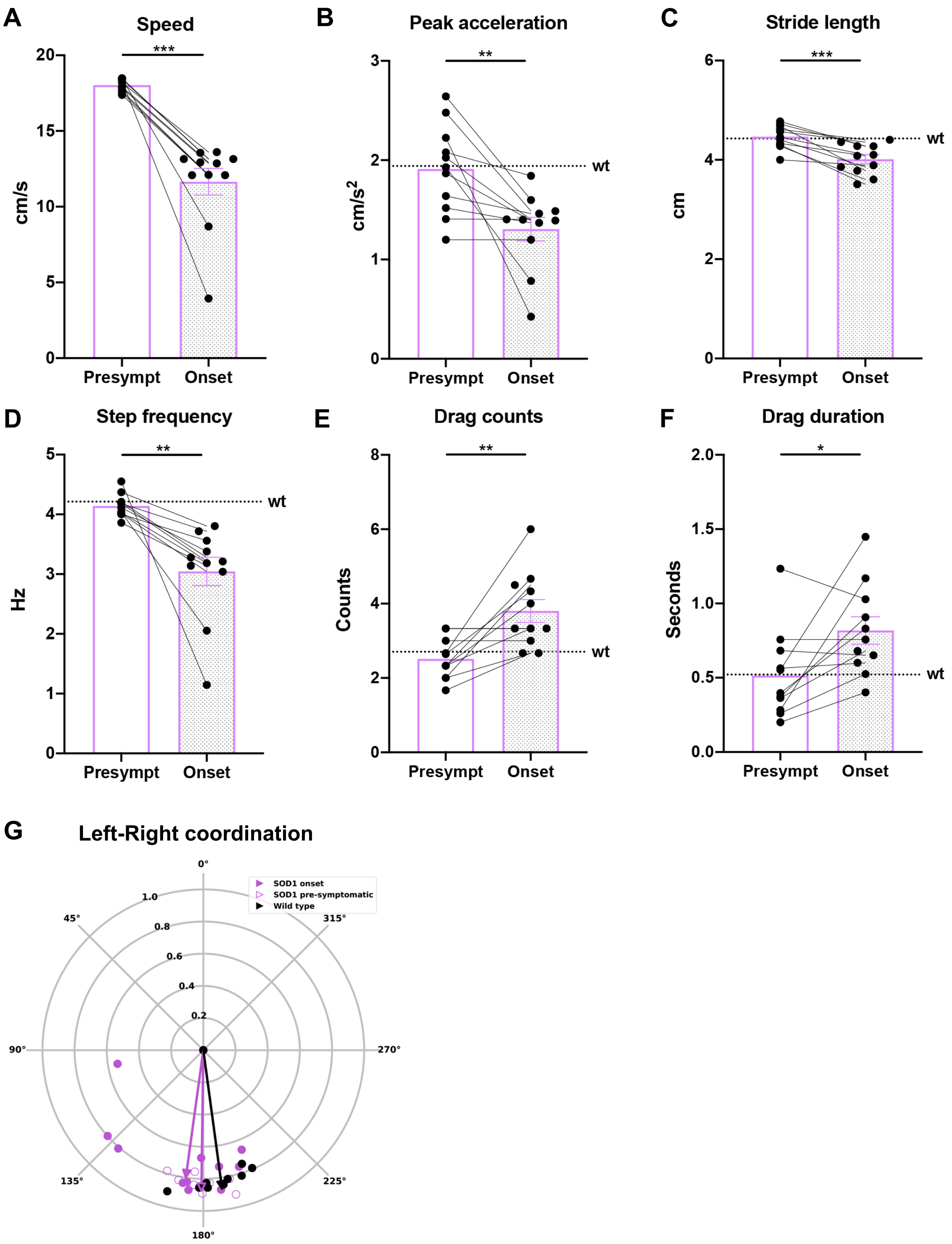


Supplementary figure 5
